# Closed-loop imitation learning reveals muscle-centric and latent-goal codes in primate sensorimotor cortex

**DOI:** 10.64898/2026.02.01.703133

**Authors:** Alessandro Marin Vargas, Adriana Perez Rotondo, Alberto Silvio Chiappa, Mackenzie Weygandt Mathis, Alexander Mathis

## Abstract

Dexterous grasping requires the seamless integration of proprioceptive feedback with predictive motor commands. Yet, how cortical circuits combine afferent feedback with efference copies to support skilled hand control remains poorly understood. Here we develop a closed-loop, muscle-level model of primate grasping that integrates biomechanics, imitation learning, and neural recordings. A neural network policy trained on a 39-muscle musculoskeletal hand reproduces naturalistic pre-contact shaping and develops internal states that quantitatively explain single-neuron activity in primary motor (M1) and somatosensory (S1) cortices. Three principles emerged. First, muscle-based controllers generate representations that align more closely with cortical dynamics than joint-based controllers, despite lower kinematic accuracy. Second, recurrent architectures with temporal memory, especially LSTMs, provide an inductive bias that enhances neural predictability. Third, model-to-brain alignment peaked at the layer integrating proprioceptive and goal signals. Finally, by decoding the model’s latent trajectory representation from M1, we demonstrated direct neural control of the policy: with activity from only tens of neurons, the brain-driven controller generated coherent grasp trajectories and showed markedly greater robustness to noise than joint-angle decoding. These findings reveal that S1 and M1 embed integrated, temporally structured, muscle-centric states and establish a stimulus-computable mechanistic framework for modeling sensorimotor control, while opening a novel route for creating brain-body models.

## Introduction

Dexterous hand use depends on the brain’s ability to couple intended actions with the evolving mechanical state of the limb. This process relies on internal control states that coordinate many muscles, predict the sensory consequences of motor commands, and update continuously through proprioceptive feedback (1, 2), supporting the robustness and adaptability of natural movement (3–5). Primary motor (M1) and somatosensory (S1) cortices occupy a central position in this control loop, interacting with parietal, premotor, and visual areas to transform goals and afferent signals into descending muscle commands (6–8). Although neuronal tuning, somatotopy, and population dynamics in these regions have been extensively investigated (9–12), the structure of the underlying control state, what variables it encodes and how efferent and afferent signals are combined during real-time hand control, remains unclear.

Grasping provides a particularly informative context in which to address this question. Prior to contact, the hand must be preshaped using vision and proprioception alone (6, 11), requiring cortical circuits to integrate predictive signals, afferent feedback, and limb biomechanics to generate object-specific postures. Population activity related to grasping has been reported across parietal, premotor, M1, and S1 cortices (13–15), yet interpreting these dynamics remains challenging. Biomechanical constraints, sensory inflow, and task goals are tightly entangled, and neural recordings alone do not reveal how these factors jointly give rise to a control state capable of guiding behavior.

This challenge is closely tied to a longstanding question in sensorimotor neuroscience: the level of representation used by cortex to support dexterous control. Classical frameworks describe M1 and S1 activity in terms of joint angles, end-point kinematics, or other abstract variables (16–19). However, proprioceptive signals originate in muscles, tendons, and skin, and motor commands ultimately act on muscles rather than joints. Proprioception further reflects a dynamic interaction between afferent signals from muscle spindles and Golgi tendon organs and reafferent signals shaped by descending motor commands (1, 2, 20). Recent work in the mouse forelimb shows that cortical populations preferentially encode muscle-level variables and that a musculoskeletal optimal-feedback-control model can reproduce this structure (21). Whether a similar muscle-centric, feedback-integrating control state underlies primate hand control, and how it is reflected in M1 and S1 population dynamics, remains unknown.

Advances in musculoskeletal modeling and learning-based control now make it possible to interrogate internal control strategies underlying dexterous hand behavior. Biomechanical simulations combined with reinforcement learning have produced controllers capable of complex grasping and manipulation (22–25), but these task-optimized policies often diverge from natural movement statistics and rely on unrealistic sensory assumptions (25, 26). In parallel, neural network models trained on behavioral or neural data (20, 27–33) reproduce selected aspects of cortical population activity, yet typically operate in open loop or abstract away musculoskeletal structure and proprioceptive feedback. By contrast, imitation learning produces naturalistic movement statistics (34–36) while preserving physiologically realistic sensory inputs, offering a promising route to models that can bridge biomechanics, control, and neural population dynamics.

Here we introduce a computational framework that leverages this convergence to identify the internal control state that emerges when a controller is constrained by primate hand biomechanics, proprioception, and natural grasping behavior, and to test whether this state aligns with population activity in M1 and S1. Using a 39-muscle musculoskeletal hand model, we train neural network controllers via imitation learning on naturalistic pre-contact trajectories (12). These controllers operate in closed loop, generate muscle-level commands, and expose internal dynamics that can be directly compared with simultaneous cortical recordings.

By systematically varying the level of representation (muscle versus joint), network architecture, and the locus of sensory-goal integration, we uncover three organizational principles. Muscle-based controllers develop internal dynamics that more closely match M1 and S1 population activity than joint-based controllers, even when joint-based models better predict overt kinematics. Recurrent architectures with temporal memory, particularly LSTMs, provide a critical inductive bias for capturing the evolving, history-dependent structure of cortical activity. Model-brain alignment is strongest at the layer where proprioceptive and goal-related signals are integrated, consistent with cortical populations encoding an abstract control state rather than raw sensory or motor variables. Decoding this latent state from M1 further enables real-time control of the model, yielding stable, object-consistent grasps with only tens of neurons and improved robustness compared to joint-angle decoding.

Taken together, these results support the view that primate M1 and S1 express a temporally structured, muscle-centric internal state that integrates proprioceptive feedback with movement goals to guide dexterous hand actions. Our framework provides a stimulus-computable, biomechanically grounded account of sensorimotor control and establishes a principled link between cortical population dynamics and the physical variables governing real hand movement.

## Results

### Neural population structure during object-specific pre-shaping

A central goal of this study is to identify the computational principles that link cortical population activity to the control of dexterous movement. To anchor our modeling framework in biological data, we first examined how sensorimotor cortex represents hand configuration during object-specific pre-shaping, the phase of grasping prior to contact, dominated by predictive and proprioceptive signals.

Primate grasping relies on a feedback loop that integrates visual input, parietal planning, premotor coordination, and sensorimotor processing in M1 and S1 (Fig. 1A) (11–13). To examine how these circuits represent anticipatory hand configurations, we analyzed a dataset in which macaques grasped 35 objects of varying shapes, sizes, and orientations while single-unit activity was recorded simultaneously in M1 and S1 (12). We focused on the pre-contact window, from movement onset to object touch, when proprioceptive and motor signals configure the hand in anticipation of grasp (Fig. 1B). This period isolates cortical activity related to internal estimates of hand state (14, 15, 37). The dataset included multiple non-human primates, sessions, and neurons from both cortical areas (Supp. Fig. S1A).

**Fig. 1.**
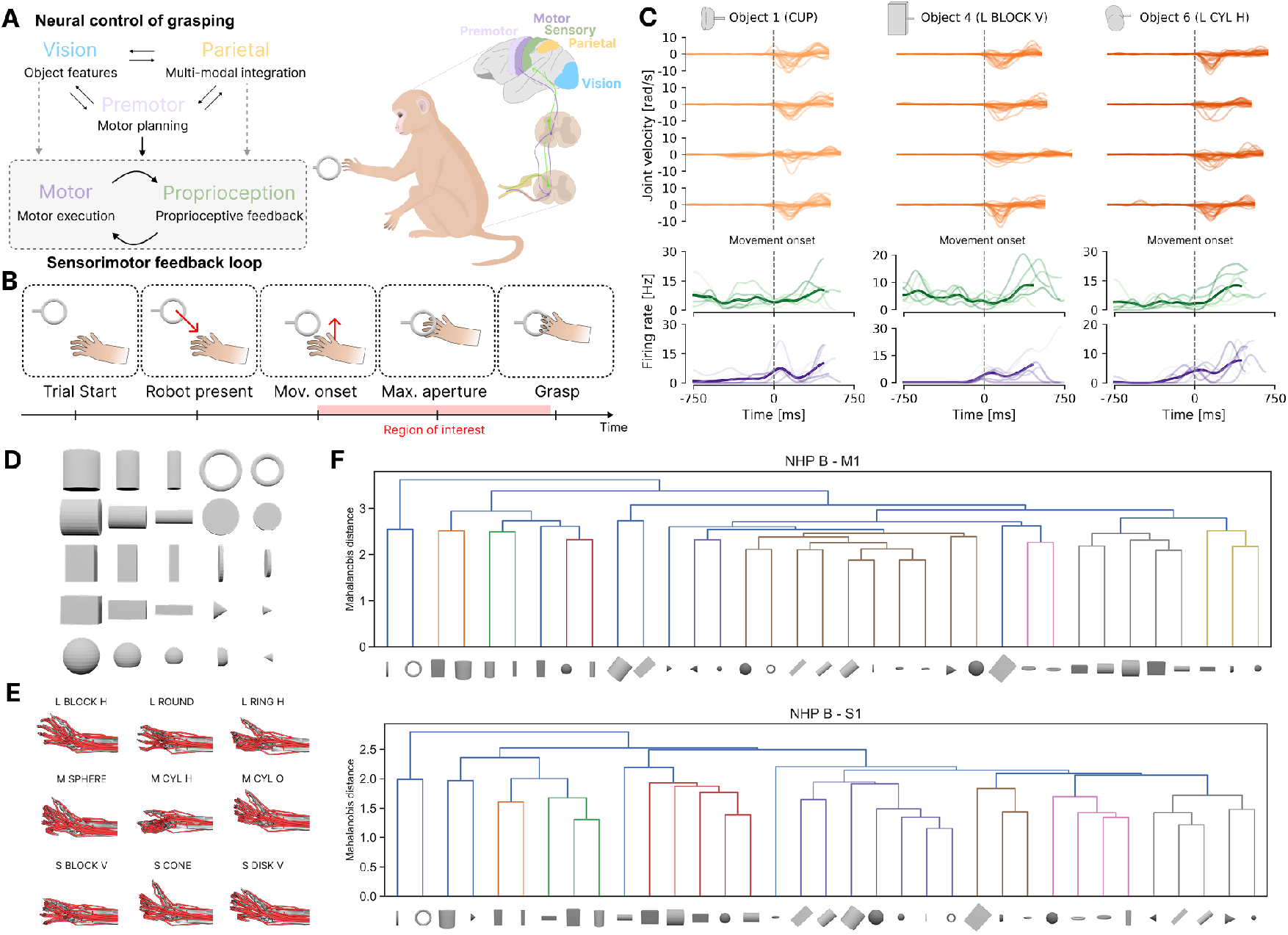
Behavioral dataset and neural representational structure. **A**. Schematic of neural control of grasping highlighting the sensorimotor feedback loop linking visual, parietal, premotor, motor, and proprioceptive pathways. **B**. Trial timeline from start to grasp; the red bar marks the pre-contact (pre-shaping) analysis window. **C**. Example joint velocities (top) and single-unit firing rates from S1 (middle) and M1 (bottom) aligned to movement onset for three different objects, illustrating across-trial variability. **D**. Set of objects grasped in the behavioral task. **E**. Example musculoskeletal hand postures at the end of the pre-shaping period for several objects, showing object-specific configurations. **F**. Hierarchical clustering of neural population activity across objects. In both M1 (top) and S1 (bottom), objects with similar orientations or sizes group together, revealing representational similarity structures that reflect task-relevant object properties.

Across sessions, both kinematics and neural activity varied systematically with object identity. Example joint velocities and single-neuron firing rates revealed object-specific temporal structure and trial-to-trial variability (Fig. 1C-E). Representational dissimilarity matrices (RDMs) computed across objects showed that both M1 and S1 were organized by physical object properties such as orientation and size (Fig. 1F). As a control, hierarchical clustering of object-specific joint configurations revealed a similarly graded organization based on geometry (Supp. Fig. S1B), confirming that both behavioral and neural representations share a common low-dimensional structure but leaving open whether cortical encoding merely reflects kinematics or abstracts them into higher-level features.

Before developing closed-loop models, we established a quantitative baseline to assess how well simple biomechanical variables explain cortical activity. To capture muscle-level biomechanics, we derived two sets of regressors (Supp. Fig. S1C): *passive* features obtained via inverse kinematics (IK), muscle length and velocity, and *active* features obtained via inverse dynamics (ID), which additionally estimate muscle activation and force (See Methods). These variables serve as biological proxies by approximating proprioceptive and reafferent inputs available to S1 and M1 (1–4).

Regression models were trained on 70% of trials, validated on 10%, and tested on 20% held-out IID trials, as well as on objects entirely excluded from both policy and regression training (OOD). Unless otherwise stated, results are reported on this strict OOD test. Linear ridge regression models trained on these features captured a meaningful fraction of neural variance in both cortical areas, with M1 neurons generally more predictable than S1 (Supp. Fig. S1D-E). However, adding active components from ID provided no systematic advantage over passive IK features, indicating that static linear combinations of muscle descriptors are insufficient to capture the integrated, feedback-driven dynamics observed in cortical populations. When each variable was tested individually, muscle length, velocity, activation, or force, none accounted for substantially higher variance than the others (Supp. Fig. S1D), reinforcing the idea that cortical activity depends on the combined state of multiple muscles states rather than any single mechanical signal.

Together, these analyses define the biological and computational constraints for the modeling framework developed in subsequent sections.

### An imitation-learning controller reproduces naturalistic pre-shaping across objects

To capture the closed-loop dynamics of proprioception, efference, and goal signals more directly, we developed a biologically inspired controller trained via imitation learning. The controller was designed to reflect a neuroanatomical hypothesis about pre-contact shaping: visual information about the object is transformed in parietal and premotor cortex into a prospective hand configuration (a *goal/trajectory* plan), while S1 conveys proprioceptive feedback and M1 integrates both proprioception and efference copies to update motor output (5, 7, 11, 13, 38) (Fig. 2A). In the model, this prospective hand configuration is provided as a short-horizon, task-conditioned reference rather than an explicitly updated plan, allowing feedback to shape control through proprioceptive and efference pathways.

**Fig. 2.**
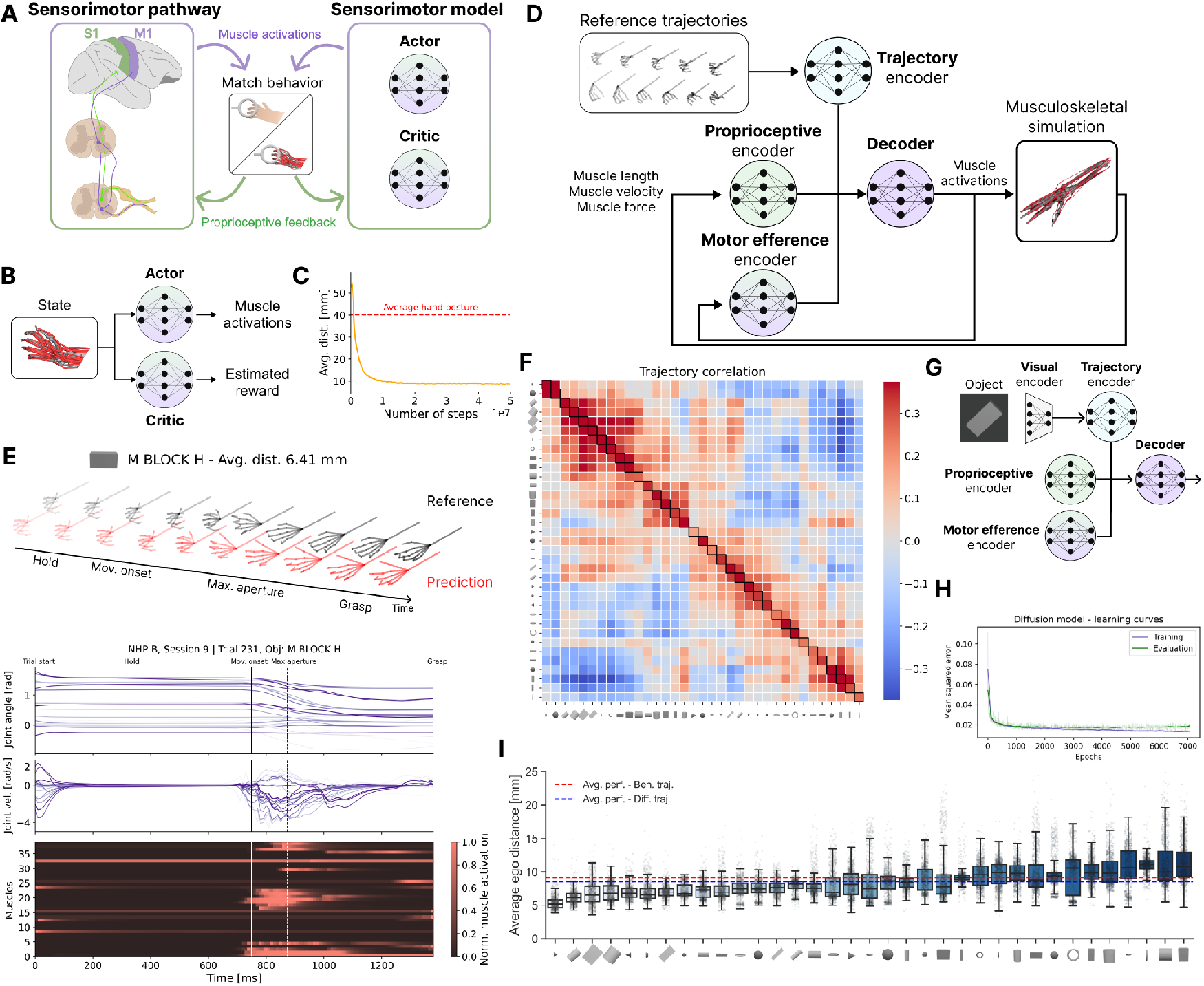
Closed-loop imitation-learning controller reproduces naturalistic grasping behavior. **A**. Conceptual link between biological sensorimotor pathways, where proprioceptive feedback and motor efference are integrated in S1/M1, and the closed-loop neural network controller. **B**. Actor-critic training scheme: the actor predicts muscle activations while the critic estimates the discounted cumulative reward. **C**. Training curve example showing the average egocentric distance as a function of training steps; the red dashed line indicates the baseline of holding the average hand posture. **D**. Policy network architecture: reference trajectories are encoded by a goal module, while proprioceptive and motor-efference streams are encoded separately; the three streams converge in a decoder that outputs muscle activations to the musculoskeletal hand model. In return, the model provides the muscle kinematics and muscle activations used as input at the next timestep. **E**. Example trial for object *M BLOCK H*, showing predicted versus reference trajectories across phases (top), joint angles and velocities (middle), and normalized muscle activations (bottom). **F**. Correlation matrix of grasping trajectories generated by the policy across all object pairs, revealing clustering by object size and orientation, consistent with behavioral organization. **G**. Vision-conditioned variant: a diffusion model maps a single egocentric object image to a kinematic trajectory, which then serves as input to the goal encoder to guide the policy. **H**. Training curve of the diffusion model showing train and validation loss. **I**. Final performance of the vision-conditioned controller across all objects: distributions of average egocentric distances, with dashed lines indicating the average imitation performance across objects between experimental and diffusion-generated trajectories.

To model these principles, the controller factorizes sensory-motor inputs into three streams that are later integrated to issue muscle activations: (1) a trajectory encoder receiving a short horizon of joint angles (10 steps, ∼ 100 ms), analogous to a short parietal/premotor goal plan (11, 29); (2) a proprioceptive encoder processing muscle length, velocity, and force, modeling afferent input to S1/M1 (1, 2); and (3) a motor-efference encoder receiving previous activations, providing a compact efference history for predictive control (3, 4). The three streams converge in a decoder that outputs muscle activations to a 39-muscle musculoskeletal hand model (22, 25, 39). The generated muscle activations update the model’s state, producing new proprioceptive feedback that is fed back into the controller, thus completing the sensorimotor loop (Fig. 2D).

Training such a controller from behavioral data presents a key challenge: expert muscle activations are not available, and the mapping from joint kinematics to activations is inherently ill-posed. Although MuJoCo provides differentiable physics, the full control problem remains non-differentiable in practice because demonstrations specify only kinematic outcomes, not the underlying muscle forces or activations. Because control operates at the muscle rather than joint level, direct supervised mapping from kinematics to activations is infeasible. This setting falls under *imitation learning from observations only* (40, 41), where demonstrations provide only kinematic outcomes. We therefore adopted a reinforcement learning approach in which the policy learns to generate muscle activations that minimize deviations from demonstrated trajectories.

Specifically, we implemented an actor-critic scheme: the actor outputs muscle activations at each timestep, while the critic estimates expected returns to guide optimization (Fig. 2B). This organization reflects both practical and biological motivations. Practically, actor-critic methods achieve state-of-the-art performance in high-dimensional musculoskeletal control (25). Biologically, the actor parallels cortical motor pathways issuing efferent commands, while the critic mirrors evaluative signals carried by basal ganglia circuits (42, 43). The reward combined penalties on joint-angle and keypoint errors with an energy cost on muscle activation, thereby encouraging movements that were accurate yet energy-efficient (See Methods).

Training converged after approximately 20 million steps, reaching stable performance well above a fixed-posture baseline (Fig. 2C). The controller reproduced pre-shaping movements across objects with an average egocentric keypoint error of 9 mm (∼13% of pinky length) (Fig. 2I; Supp. Fig. S2A). Temporal error profiles revealed distinct behavioral phases, stable hold, rapid opening, and grasp shaping, with maximal deviations during fast transitions. Predicted joint kinematics and muscle activations closely tracked recorded trajectories, showing phase-specific recruitment of muscles for stabilization, opening, and closure (Fig. 2E).

Importantly, the learned controller also captured object-level structure: correlations among predicted trajectories across objects revealed clustering by size and orientation (Fig. 2F), paralleling the organization of neural representations observed in M1 and S1 (Fig. 1F). This convergence suggests that both biological and learned systems encode grasping goals in a shared low-dimensional geometric space tied to object shape and orientation.

To probe the functional contributions of each input stream, we systematically perturbed or ablated them during both training and evaluation (Supp. Fig. S2B and See methods). Proprioceptive feedback proved essential: its removal caused the largest performance drop, and fixed or time-shifted inputs strongly degraded accuracy. Noise injections during evaluation further disrupted stability, emphasizing the importance of continuous sensory updating. The trajectory encoder was robust to moderate noise but sensitive to zeroing or temporal delays, consistent with its role in maintaining goal-directed behavior. By contrast, the previous-activation (efference) stream contributed minimally under the present task conditions, which involved stereotyped pre-shaping in the absence of external perturbations or unexpected loads. We therefore hypothesize that efference history may play a more prominent role during perturbed or uncertain interactions, where predicting the sensory consequences of motor commands becomes critical for rapid correction. Together, these manipulations indicate that proprioception and goal-related signals dominate control during unperturbed pre-shaping, while efference provides a secondary stabilizing signal whose contribution likely depends on task demands.

Finally, to verify that the controller can generalize beyond oracle trajectories, we implemented a vision-conditioned variant in which a diffusion model mapped a single egocentric image of each object to a kinematic trajectory that conditioned the policy (Fig. 2G-H; Supp. Fig. S2C-E and see methods). The diffusion model accurately reconstructed plausible pre-shaping paths and yielded controller performance comparable to that of trajectory-conditioned training (Fig. 2I). This demonstrates that the framework is compatible with visual goal inference, although all subsequent analyses focus on trajectory conditioning to isolate proprioceptive-motor integration.

Together, these results show that imitation learning from observations yields high-fidelity, biologically interpretable controllers that reproduce naturalistic pre-shaping behavior, reveal object-specific structure consistent with cortical representations, and depend critically on proprioceptive and goal inputs. This provides a grounded computational substrate for testing whether behavioral-aligned internal states align with neural population activity in M1 and S1.

### Behavioral-aligned internal states predict both single-trial variability and condition-averaged cortical responses

Having established that the imitation-trained controller reproduces naturalistic pre-shaping across objects, we next asked whether its internal states align with neural activity in sensorimotor cortex. This analysis tests a key prediction of closed-loop control theories: that the same internal variables supporting stable behavioral control should also account for both the consistent, condition-averaged patterns and the trial-to-trial fluctuations observed in cortical dynamics. Previous models of motor control often capture average tuning across conditions but do not model within-condition variability, an essential feature of population activity in M1 and S1 that reflects ongoing sensorimotor integration.

To address this, we examined whether behavioral-aligned representations could predict single-neuron firing rates across trials and conditions, and compared their performance against multiple baselines. We considered (*i*) linear encoding models built from the same experimental variables provided to the policy (muscle kinematics and joint angles), and (*ii*) untrained networks with matched architectures but random weights. For each network layer, activations during the pre-shaping window were reduced to 78 principal components (matching the dimensionality of linear baselines) and used in ridge regression to predict single-neuron firing rates. For the regression models, we used the same dataset split of the linear models.

Example neurons illustrate that trained networks capture both rapid, trial-specific fluctuations and robust, object-specific modulations. In single trials, the decoder layer predictions tracked moment-to-moment variability in S1 and M1 neurons (Fig. 3A and Supp. Fig. S3A). When responses were averaged across repetitions, the same states reproduced condition-specific tuning (Fig. 3B and Supp. Fig. S3B). This indicates that behavioral-aligned networks not only stabilize representations across repeated grasps but also retain sensitivity to trial-level dynamics inherent to cortical control.

**Fig. 3.**
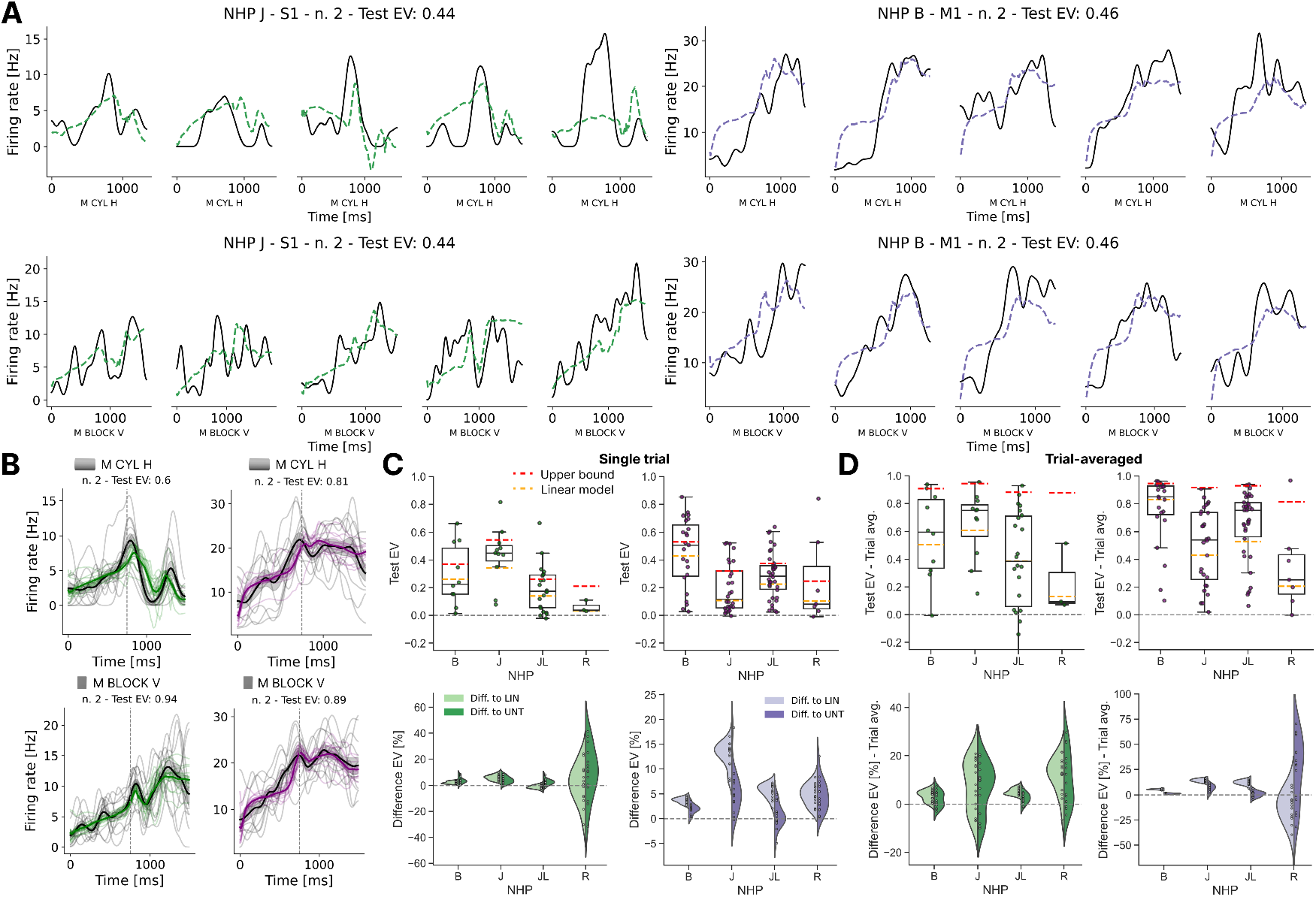
Internal states of the imitation-learning controller predict neural activity in S1 and M1. **A**. Example single-trial fits for held-out out-of-distribution (OOD) objects. Black traces: recorded firing rates from individual neurons; dashed colored traces: predictions from the decoder layer of the value network. Left, S1 neurons (green); right, M1 neurons (purple). **B**. Trial-averaged predictions for the same example neurons and objects. Thick black lines: condition-averaged firing rates; colored lines: trial-averaged predictions; thin lines: individual trial predictions; shaded areas: s.e.m. Model states capture both trial-to-trial variability and object-specific tuning. **C**. Single-trial neural predictability on OOD objects. *Top:* distributions of Test EV across neurons in each NHP, using the first decoder layer of the critic as the predictive representation. Each dot corresponds to a single neuron; orange dots show linear baseline fits using muscle kinematics; red dashed lines indicate reliability-based upper bounds (See Methods). *Bottom:* distributions of neural explainability, defined as the mean EV gain across neurons for each model relative to baselines. Here each dot corresponds to a trained network. Trained networks consistently outperform both linear regressors and untrained architectures (Mann-Whitney tests). **D**. Condition-averaged predictions. Same analyses as in **C** but using trial-averaged firing rates, showing consistent gains across NHPs and cortical areas.

Population-level analyses confirmed these findings. Using the critic’s first decoder layer as the predictive representation, we quantified the explained variance (EV) of single-neuron activity across S1 and M1. behavioral-aligned networks exhibited systematically higher EV distributions compared to both linear encoding models and untrained architectures, indicating that behavioral-aligned representations more effectively captured the variance structure of cortical population activity (Fig.3C top). To assess alignment at the model level, we computed *neural explainability*, defined as the mean EV gain relative to baselines after selecting the best predictive layer for each neuron based on training data (Fig.3C bottom). This analysis revealed consistent improvements across all trained models and animals, demonstrating that networks optimized for control outperform both static and untrained baselines in explaining neural responses. Similar effects were observed for condition-averaged predictions (Fig.3D), and alignment remained robust even when temporal offsets of up to *±* 500,ms were introduced between neural activity and network states (Supp. Fig. S3C).

As an additional control, we compared neural predictivity between the behavioral-aligned (imitation-learned) model and a purely data-driven variant trained to map muscle kinematics directly onto neural activity without motor control objectives (Supp. Fig. S3D-E). Across all NHPs, behavioral-aligned representations significantly outperformed data-driven ones in predicting cortical responses. This demonstrates that optimizing a closed-loop control objective yields internal states that are more aligned with sensorimotor cortical activity than direct regression from behavioral data alone.

Together, these results demonstrate that behavioral-aligned internal states provide a stimulus-computable account of cortical dynamics. Networks trained to reproduce dexterous grasping explain both single-trial variability and stable condition tuning in S1 and M1, outperforming static linear baselines and untrained architectures, as well as data-driven models.

### Muscle-based controllers outperform joint-based ones in neural alignment

Having shown that behavioral-aligned internal states predict both trial-specific and condition-averaged neural responses (Fig. 3), we next asked what type of control policy best gives rise to such cortical-like representations. A long-standing debate in motor neuroscience concerns whether the brain encodes hand movements in joint-centric coordinates, as often assumed in robotics and optimal feedback control models (16, 19, 48, 49), or in muscle-centric coordinates that more directly reflect proprioceptive and force signals (21, 50). This distinction is critical for understanding how afferent and efferent signals are integrated during dexterous control.

To address this question, we trained two classes of policies that differed in both their control outputs and sensory representations. *Joint-based* controllers operated in kinematic space, receiving feedback in terms of joint angles and angular velocities and issuing torques at the level of kinematic degrees of freedom. By contrast, *muscle-based* controllers received proprioceptive input in terms of muscle length, velocity, and force and generated activations for 39 individual muscles (Fig. 4A), thereby preserving the nonlinear dynamics and redundancy of the musculoskeletal system. This contrast isolates a high-level, joint-centric control representation from a muscle-centric one grounded in physiological sensing and actuation.

**Fig. 4.**
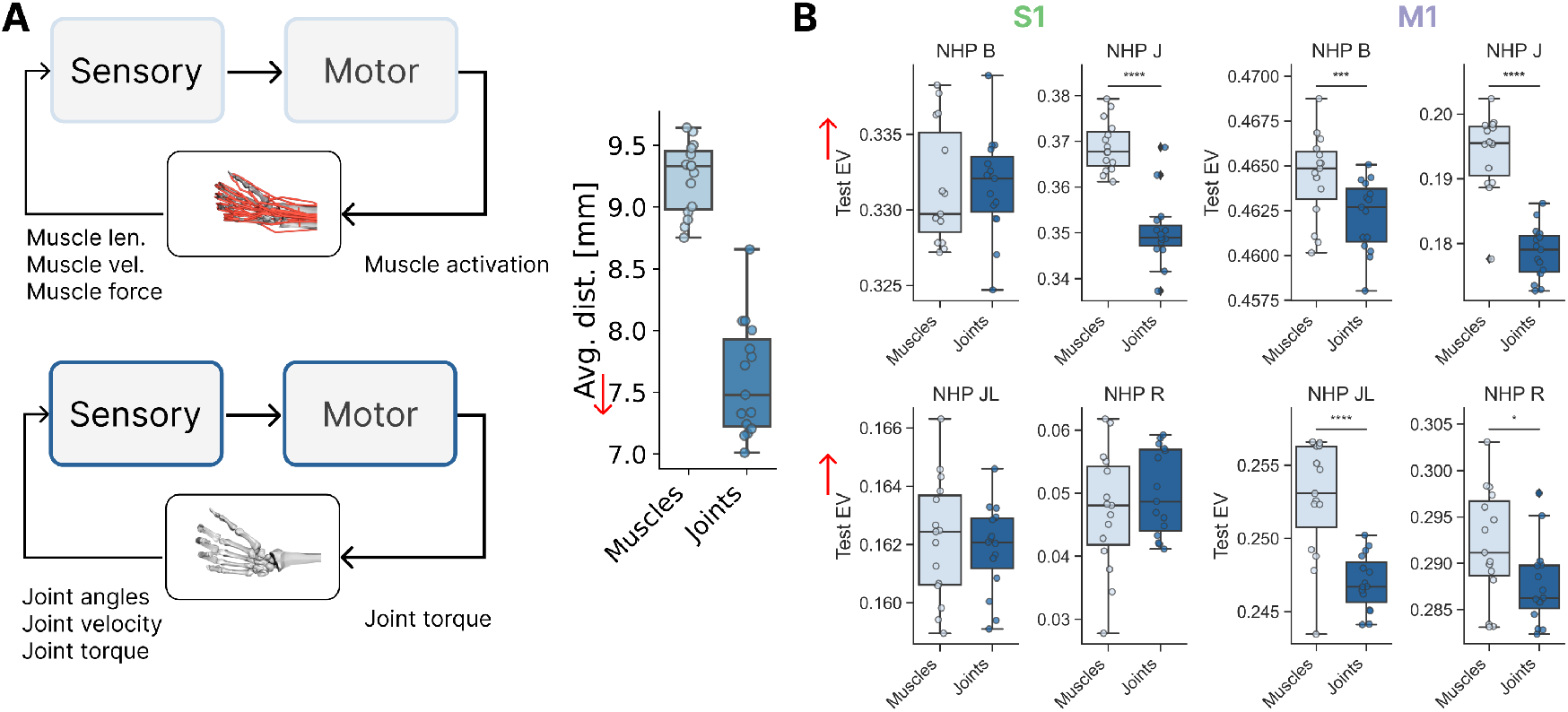
Muscle-based representation outperform joint-based ones. **A**. Schematic of control policies using muscle-based (top) versus joint-based (bottom) representations. Muscle policies output activations for 39 individual muscles, whereas joint policies output torques at the level of kinematic with 23 degrees of freedom. **B**. Behavioral performance distributions (egocentric keypoint error) show that joint-based controllers achieve lower imitation errors than muscle-based controllers. **C**. Neural explainability, defined as the mean explained variance (EV) gain relative to linear baselines, for muscle-versus joint-based controllers. Each dot corresponds to a trained network; muscle-based policies consistently yield higher neural alignment than joint-based ones, despite their lower behavioral fidelity (*: p<0.01; **: p<0.01; ***: p< 0.001, ****: p<0.0001 - Mann-Whitney tests).

As expected, joint-based policies achieved lower imitation errors and thus tracked reference trajectories more accurately (Fig. 4B). This advantage reflects the reduced dimensionality of joint control, which bypasses muscle nonlinearities and is commonly exploited in engineered systems (34, 39). However, when we examined neural alignment, the opposite pattern emerged: muscle-based policies consistently outperformed joint-based ones, with the effect being robust and statistically significant in M1, but not in S1 (Fig. 4C). The nonsignificant trend in S1 suggests that its encoding of proprioceptive signals may be captured by both joint- and muscle-level models, whereas M1 preferentially reflects muscle-based control variables.

To rule out the possibility that this effect arose solely from the input features, we compared linear encoding models built on muscle versus joint kinematics (Supp. Fig. S4). These baselines performed similarly, indicating that the advantage of muscle-based controllers does not stem from trivial feature differences but from the representational structure that emerges under closed-loop muscle-level optimization.

Together, these results reveal a dissociation between behavioral and neural criteria: while joint-based control maximizes task accuracy, muscle-based control produces internal states that align more closely with cortical activity, particularly in M1. This finding extends physiological evidence that motor cortex is more strongly coupled to muscle-level variables than to joint kinematics (21), and highlights potential differences in how M1 and S1 contribute to the integration of proprioceptive and motor signals.

### Sequential temporal processing boosts neural predictability

Having shown that muscle-based controllers best align with cortical activity, we next asked which computational mechanisms within a control architecture promote this alignment. In sensory neuroscience, task-optimized convolutional networks have provided a unifying framework linking architecture and function, showing that ventral stream responses can be predicted from hierarchically structured visual models (5, 51, 52).By analogy, we tested a set of biologically motivated hypotheses about how proprioceptive and motor signals might be organized and integrated during dexterous control (Fig. 5A and Table 1).

**Table 1.** Architectural hypotheses tested. Encoder/Decoder specify how proprioceptive, goal, and efference streams are processed and merged. Abbrev.: GCN = Graph Conv. Net; GAT = Graph Attention Net; TCN_1_ = channel-wise temporal conv.; modLSTM = per-stream LSTM.

| Encoder | Decoder | Integration focus | Hypothesis / biological analogy |
| --- | --- | --- | --- |
| MLP | MLP | Global all-to-all (spatial) | Distributed muscle mixing without hard anatomical constraints; broad population coupling in cortex (10, 12). |
| GCN / GAT | MLP | Local structured (spatial) | Fixed or attention-weighted muscle connectivity; partial modularity / spinal-like synergies (44, 45). |
| Transformer (enc) | MLP | Dynamic context (spatial) | Context-dependent reweighting of muscle features with simple readout; flexible cortical coupling (46, 47). |
| Transformer (enc) | Transformer (dec) | Dynamic context (end-to-end) | Fully attentional integration and readout; hypothesis of maximally flexible coordination. |
| TCN | MLP | Short-horizon spatiotemporal filter | Fixed-window temporal dependencies over all channels; short-latency feedback loops. |
| TCN<sub>1</sub> | MLP | Per-channel temporal filter | Shared short-term filtering per proprioceptive channel before fusion. |
| MLP | LSTM | Adaptive temporal (global) | Recurrent latent state maintains and updates context for predictive control (27, 29). |
| TCN / TCN<sub>1</sub> | LSTM | Short + adaptive temporal | Local temporal filtering feeding a recurrent state; multiscale temporal control. |
| LSTM (proprio-only) | LSTM | Modular recurrent | Proprio stream recurrently encoded; other streams via MLP; recurrent decoder integrates for multi-muscle feedback. |
| modLSTM (per stream) | LSTM | Hierarchical recurrent | Each stream is recurrent (per-stream LSTM); recurrent decoder integrates stream-specific memories (distributed gating). |

**Fig. 5.**
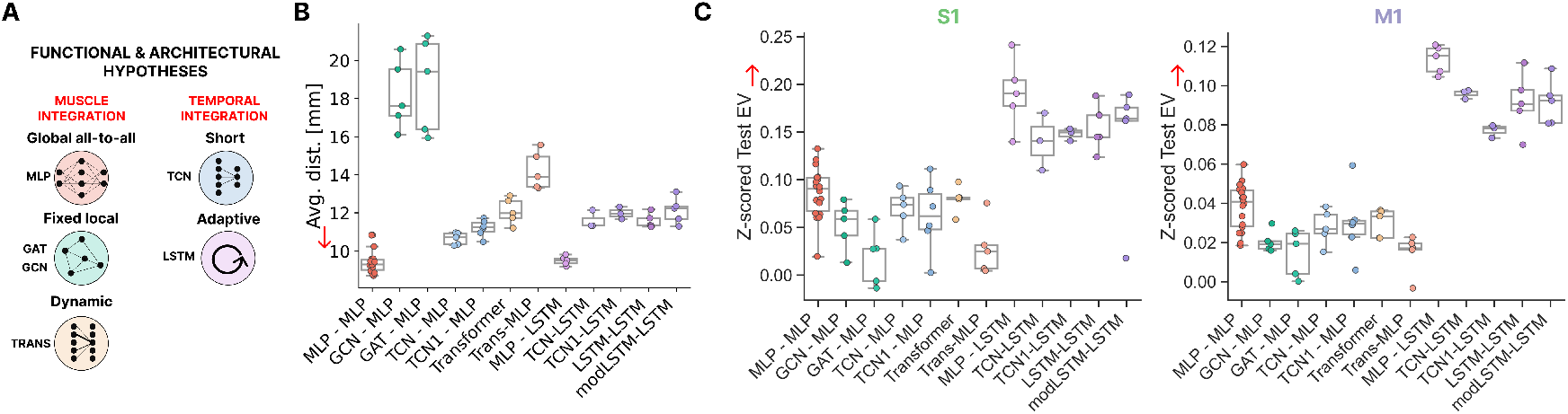
Functional hypotheses tested through architectural comparisons. **A**. Schematic linking architectural choices to functional hypotheses about sensorimotor integration. Left: *muscle integration*, comparing global all-to-all (MLP), fixed local (GCN/GAT), and dynamic (Transformer) connectivity. Right: *temporal integration*, contrasting short-range (TCN) versus adaptive recurrent (LSTM, modLSTM) mechanisms. **B**. Task performance across architectures, measured by average egocentric distance (lower is better). Each dot represents a trained network. **C**. Neural alignment across architectures, quantified as z-scored explained variance (EV) relative to within-animal baselines, allowing pooling across NHPs. Results are shown separately for S1 (left, green) and M1 (right, purple). Each dot corresponds to a single trained network. Architectures with recurrent or adaptive temporal integration (LSTM, modLSTM) yield consistently higher neural predictability.

We considered two broad functional dimensions (Fig. 5A): (i) *muscle integration*, how proprioceptive inputs from multiple muscles are combined, and (ii) *temporal integration*, how current motor commands incorporate past sensory history. For spatial integration, we compared three network architectures. First, a global all-to-all scheme (MLP) in which proprioceptive and efferent features interact without anatomical constraints, akin to dense coupling observed across muscles in M1 population codes (10, 12, 53). Second, a fixed local scheme (GCN, GAT) in which muscles interact through a predefined or attention-weighted neighborhood graph, motivated by evidence for partial modularity and muscle-synergy structure in spinal circuits (44, 45). Third, a dynamic integration scheme (Transformer) in which attention weights are updated at each timestep, reflecting the flexible context-dependent coupling between cortical neurons during grasp (46, 47). For temporal integration, we contrasted short, feedforward timescales (Temporal Convolutional Network, TCN) with adaptive, recurrent architectures (LSTM, modLSTM) that maintain internal states over time. These correspond to alternative hypotheses about how motor cortex encodes temporal context: either as a fixed filter of recent sensory inputs or as an actively maintained latent trajectory that supports predictive control (27, 29, 31). Recurrent mechanisms, in particular, have been proposed to underlie the “dynamical systems” perspective of M1 activity (54, 55), allowing population activity to evolve continuously through internal feedback loops.

We trained each architecture as a closed-loop controller under identical behavioral conditions and compared both task performance and neural alignment (Fig. 5B-C). Behaviorally, MLP and LSTM architectures achieved the lowest imitation errors, followed by TCN and Transformer variants, while graph-based models (GCN, GAT) performed worst. The latter’s limited flexibility likely reflects their fixed spatial constraints, which may be more relevant to spinal-level coordination than to cortical control.

Neural analyses revealed a clear pattern: architectures that maintained an internal temporal state, particularly LSTM-based and modular recurrent models, consistently achieved higher explained variance across neurons, sessions, and animals (Fig. 5C). In contrast, models lacking recurrence (MLP, TCN, Transformer) primarily captured static or short-timescale dependencies and were less able to predict the evolving structure of population activity in M1 and S1. These results were robust to the choice of activation extraction method (see Methods; Supp. Fig. S5A). Importantly, neural predictability was not strongly correlated with task performance, indicating that architectural differences, rather than behavioral accuracy alone, primarily accounted for differences in neural alignment (Supp. Fig. S5B-C).

Together, these results do not imply that sensorimotor cortex implements a specific recurrent architecture. Rather, they suggest that sequential temporal integration, the ability to retain and update internal state over behaviorally relevant timescales, is a key organizational principle for reproducing cortical population dynamics during grasp. Incorporating such temporal memory provides a biologically grounded inductive bias for modeling closed-loop control, helping bridge the gap between behavioral competence and neural representational structure.

### Sensorimotor representations are best explained by the convergence of sensory and motor information

The preceding analyses identified temporal memory as a critical inductive bias for aligning control policies with cortical population activity. We next asked where within the trained controller such alignment with cortical activity emerges, whether in layers dominated by sensory or motor processing, or in those integrating the two. This question addresses a long-standing issue in motor neuroscience: whether cortical populations maintain segregated representations of proprioceptive and efferent variables or instead form unified sensorimotor states that jointly encode feedback and intention (4, 12). Physiological evidence indicates that both S1 and M1 receive converging afferent and efferent signals during grasp and manipulation (7, 13), yet the degree to which these signals are fused within cortical circuits remains unresolved.

Before testing the model against neural recordings, we first performed a synthetic control experiment to interpret how network modules map onto one another. Using a “student-teacher” framework, a simplified controller (“Pose Policy”) was trained to maintain a fixed grasp posture, producing deterministic muscle activations and joint kinematics. A second network (“Student Policy”) was then trained to imitate the teacher’s behavior using only observed kinematics, without access to internal states (Supp. Fig. S6A-B). We then asked whether principal components (PCs) from individual modules of the student could predict those from the corresponding modules in the teacher. As expected, each student encoder best predicted its homologous module in the teacher, confirming a clear one-to-one correspondence across network hierarchies. Nonetheless, other encoders also captured a substantial fraction of variance, reflecting partial overlap between representations. The decoder modules, responsible for integrating sensory and motor streams, showed the highest overall mutual predictability (Supp. Fig. S6C), consistent with the emergence of shared latent codes that combine proprioceptive and motor information even in this simplified control task.

We next applied the same analysis framework to neural data, comparing the predictive power of each network module against single-neuron activity in S1 and M1 across four NHPs (Fig. 6A). Across individuals, predictive profiles varied but revealed a consistent pattern: while the trajectory and proprioceptive encoders explained subsets of neural variance, the decoder, where sensory and motor streams converge, yielded the highest explained variance (EV) for both cortical areas. This integrative stage combines proprioceptive feedback with motor-goal information, paralleling the convergence of ascending and descending signals in the cortical sensorimotor loop.

**Fig. 6.**
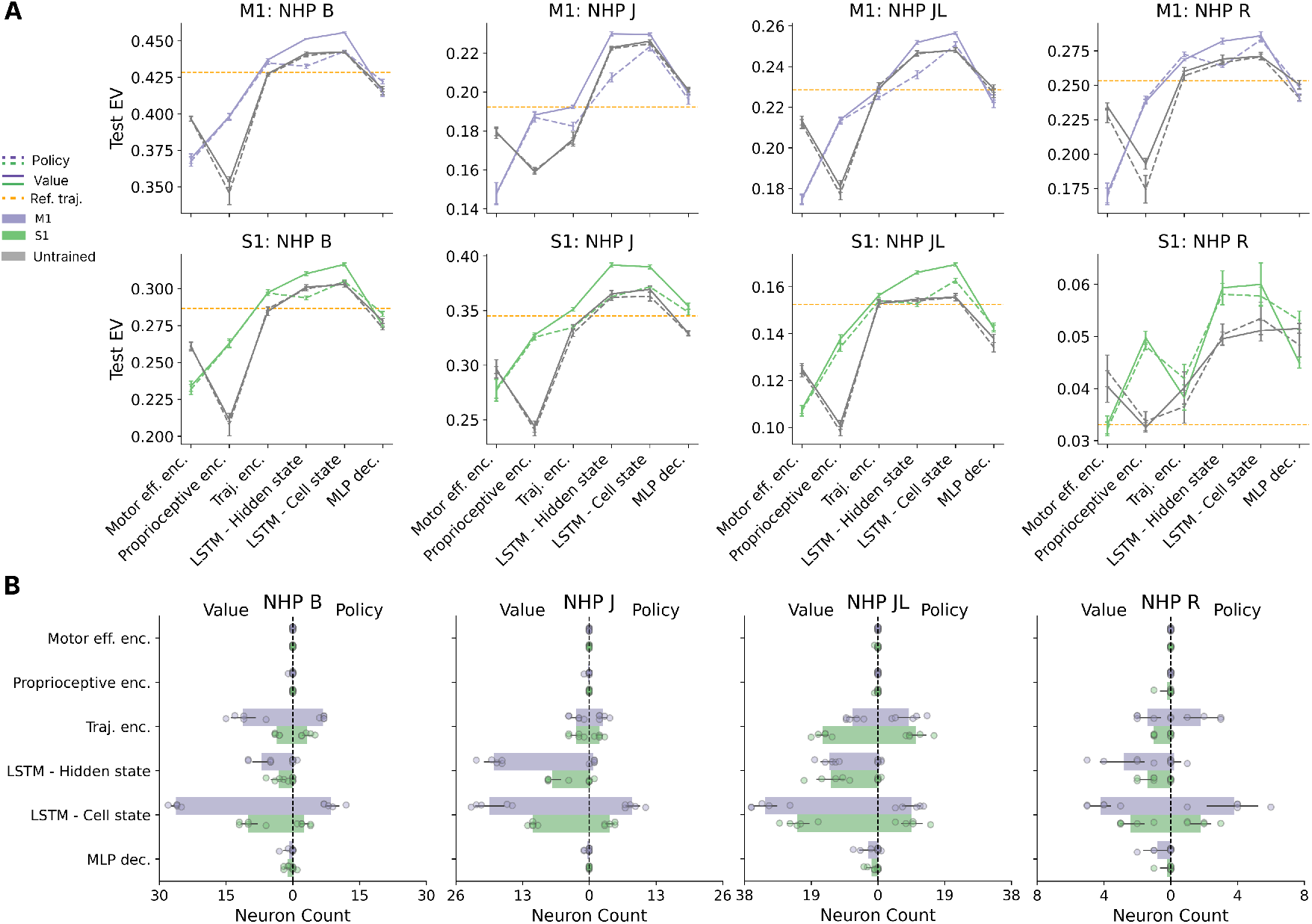
Hierarchical correspondence between network layers and cortical areas. **A**. Explained variance (EV) of neural predictions across successive layers of the trained LSTM controller for each NHP, shown separately for M1 (top, purple) and S1 (bottom, green). Each curve shows mean EV across neurons, comparing trained and untrained networks. The orange dashed line indicates the predictions achieved using the reference trajectory as baseline. Error bars denote 95% confidence intervals across five independently trained models. **B**. Distribution of preferred layers across neurons for policy and value networks. Each dot represents one trained LSTM model; horizontal bars show mean with s.e.m. across models.

To further assess how task optimization shapes these correspondences, we compared trained models with their untrained counterparts (Fig. 6A). The trajectory encoder showed minimal improvement with training, unsurprising, since the reference trajectory itself provides strong predictive information even for random networks. The actuator encoder (processing previous muscle activations) showed similarly limited gains, consistent with its weaker contribution to task performance (Supp. Fig. S2) and the underdetermined nature of muscle inverse dynamics. By contrast, both the proprioceptive encoder and decoder exhibited substantial increases in neural predictability after training, indicating that proprioceptive feedback and its integration with efferent signals are the crucial parts of neural networks representational alignment.

We also compared two extraction regimes: (i) a *dynamic* configuration, in which the network generated its own proprioceptive feedback in closed loop, and (ii) a *clamped* configuration, where inputs were replaced by measured kinematics (Supp. Fig. S6D). Both yielded comparable neural predictability, confirming that autonomous feedback dynamics capture biologically meaningful computations. However, clamping activations markedly improved performance for the policy network, particularly within the proprioceptive encoder and decoder, indicating that better imitation of behavioral trajectories strengthens neural alignment. This result suggests that the fidelity of sensory feedback, rather than its source (real or simulated), constrains the ceiling of neural predictability.

To test whether the improved neural alignment in the decoder arises from genuine integration or from a simple aggregation of input features, we compared its representations with those obtained by concatenating the outputs of the individual proprioceptive, goal, and motor efference encoders (Supp. Fig. S6E). Despite controlling for dimensionality through PCA, the integrated decoder representation consistently outperformed the concatenated one across monkeys and cortical areas. This result indicates that cortical-like alignment depends on nonlinear integration of sensory and motor streams, rather than on their additive combination, underscoring the functional importance of sensorimotor convergence for representing hand state.

Together, these results demonstrate that cortical activity in S1 and M1 is best explained by the convergence of sensory and motor streams rather than by isolated feature channels. Integration at the network’s decoder layer mirrors the architecture of cortical sensorimotor loops, in which feedback and feedforward pathways are dynamically combined to guide dexterous hand control.

### Sensorimotor representations reflect evaluation of kinematic states

The dual actor-critic architecture of the controller provides a means to ask whether cortical activity is better explained by motor command generation or by evaluation of the ongoing kinematic state. In reinforcement learning, the *policy* network produces muscle activations that directly drive the musculoskeletal system, whereas the *value* network estimates the cumulative discounted reward, an implicit measure of how favorable a configuration is for achieving the goal. This distinction parallels that between execution and evaluation processes in biological motor control, with the former linked to cortical output pathways and the latter to evaluative or predictive computations in cortico-basal ganglia circuits (4, 42, 43). We therefore examined whether neuronal activity in S1 and M1 aligns more closely with representations arising from motor generation or from state evaluation.

Across all NHPs, neural predictions based on the *value* network consistently outperformed those derived from the *policy* network in both M1 and S1 (Fig. 6A-B). This advantage was robust across layers and animals, indicating that cortical activity aligns more closely with representations reflecting the evaluation of the current kinematic state than with those directly generating motor commands. The higher explained variance (EV) of the value network suggests that its representations capture integrated sensory-motor features that jointly encode the movement goal and the quality of the current configuration. In contrast, the policy network, optimized for action generation, showed weaker alignment under dynamic feedback conditions, likely because small sensory discrepancies propagate through its closed-loop inputs. However, under clamped conditions where sensory feedback was perfectly matched to the behavioral trajectory, policy representations substantially improved, achieving prediction accuracy comparable to the value network (Supp fig. S6). This convergence indicates that both networks capture relevant aspects of cortical computation: value representations provide a stable, goal-centered evaluation of state, whereas policy representations reflect the underlying motor control signals when sensory feedback is accurate.

Among input streams, the *trajectory encoder* emerged as the most predictive component across monkeys and areas, outperforming proprioceptive and efference encoders. This result suggests that cortical populations encode a latent representation of the current and intended movement trajectory, a hypothesis directly tested in the next section.

### Brain-controlled policy

Having established that cortical activity aligns with the controller’s internal representations, we next asked whether this correspondence could be reversed. Specifically, can population activity in S1 and M1 directly *drive* the controller, replacing the reference trajectory with a brain-derived signal? Demonstrating such control would provide a causal link between cortical population dynamics and embodied motor behavior through a learned sensorimotor policy.

We implemented a *brain-controlled policy* in which population activity from S1 or M1 replaced the reference trajectory input to the controller (Fig. 7A). To test alternative hypotheses about how cortical activity can drive movement, we trained two linear decoders on neural recordings collected during natural grasping (Fig. 7B). The first, a *joint-angle decoder*, mapped neural activity directly onto joint kinematics, producing predicted poses that were applied to the musculoskeletal model without involving the learned policy. This approach corresponds to the classical paradigm in brain-machine interfaces (BMIs), where neural signals are decoded into explicit movement variables that directly drive an effector. The second, a *latent-trajectory decoder*, instead mapped neural activity onto the low-dimensional trajectory embedding learned by the policy’s encoder. In this configuration, the decoded latent features replaced the policy’s reference input, allowing cortical activity to drive the closed-loop controller through its learned internal representation. This design allowed us to determine whether cortical population dynamics naturally align with explicit kinematic variables or with the higher-level trajectory representations that the model uses to generate coordinated muscle control.

**Fig. 7.**
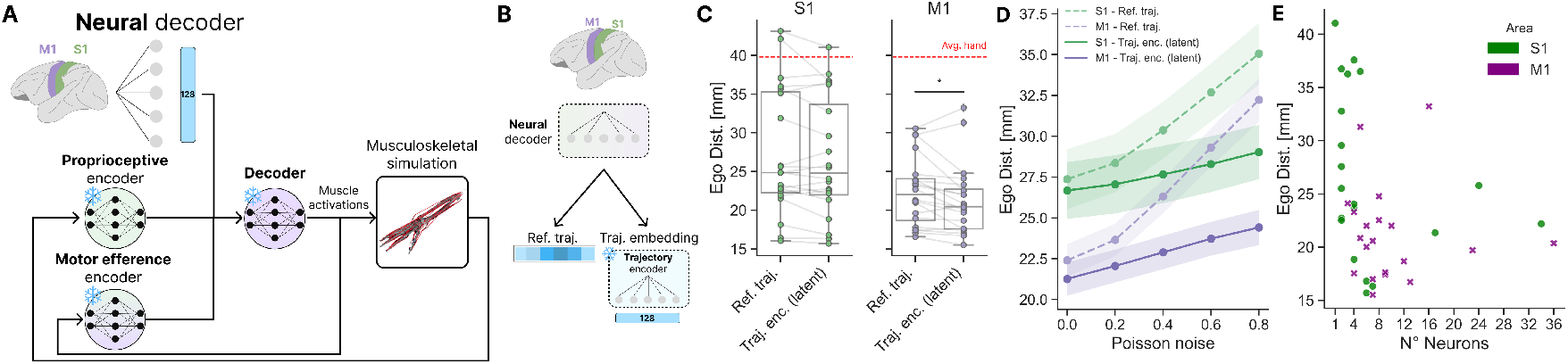
Neural activity can drive the policy through decoded trajectory representations. **A**. Schematic of the neural-controlled policy. Neural population activity from S1 or M1 is linearly decoded into latent trajectory features, which replace the reference input to the closed-loop controller. The decoder uses population firing rates to predict the corresponding trajectory embedding used by the trained policy to generate muscle activations. **B**. Two neural decoder variants. In the first (top), neural activity predicts the reference trajectory directly. In the second (bottom), neural activity predicts the latent trajectory embedding from the policy’s internal representation, allowing neural activity to interface with the controller at a representational rather than kinematic level. **C**. Behavioral performance of the brain-controlled policy. Each dot corresponds to a session, showing the egocentric distance between trajectories generated under neural control and the reference trajectories. Dashed horizontal lines indicate baseline errors from direct joint-angle decoding (green for S1, purple for M1); the red dashed line marks the error from maintaining the average hand posture. **D**. Robustness of neural decoding to Poisson-scaled noise. The curves show the mean egocentric error as a function of added neural noise for latent goal decoding (solid lines) and direct joint-angle decoding (dashed lines) from S1 (green) and M1 (purple); shaded regions indicate the standard error across sessions. Decoding trajectory embeddings yields substantially greater noise tolerance, suggesting that the learned latent representations capture more stable and robust control features than direct kinematic decoding. **E**. Relationship between decoding performance (egocentric distance) and the number of recorded neurons. Each point represents a distinct subset of neurons from S1 (green) or M1 (purple). Effective control can be achieved with relatively small neural populations, consistent with low-dimensional population dynamics underlying grasp control.

Neural activity from both S1 and M1 reliably reproduced movement-related variables (Supp. Fig. S7A), and when used to drive the controller, generated realistic grasping trajectories that closely matched behavioral data (Fig. 7C). Decoding into the policy’s latent trajectory space produced movements with comparable imitation accuracy to those obtained by direct joint-angle decoding, on average 20 mm for M1-driven control and 25 mm for S1-driven control, both well below the fixed-posture baseline. This demonstrates that the model’s learned latent representations form a behaviorally meaningful control manifold capable of supporting naturalistic movement generation. The ability to substitute cortical activity for sensory reference inputs thus provides a direct functional validation of the model, showing that its internal goal representations are not merely abstract encodings but actionable variables through which neural population activity can drive closed-loop motor behavior.

Extending this analysis, we decoded neural activity into latent representations from other modules of the controller (Supp. Fig. S7B). While all decoded layers supported above-baseline control, decoding the trajectory-goal representation from M1 yielded the lowest errors, followed by the integrative layer, whereas S1 showed relatively stronger alignment with proprioceptive latents despite lower overall control fidelity.

We next examined the stability of these neural interfaces under degraded conditions by injecting Poisson-scaled noise into the recorded activity (Fig. 7D). Although performance of both decoders declined gradually with increasing noise, the latent-trajectory decoder remained substantially more robust than direct joint-angle decoding for both S1 and M1. This robustness indicates that the policy’s latent representation captures stable, noise-tolerant features of movement, properties that mirror the smooth, low-dimensional organization observed in cortical population dynamics.

Consistent with their distinct functional roles, M1-driven decoders achieved higher control fidelity than those based on S1, reflecting M1’s direct involvement in generating motor commands. S1 decoders produced weaker but still above-baseline control, in line with its role in representing proprioceptive feedback and integrating sensory context with efferent signals (20). Effective control was obtained with only tens of simultaneously recorded neurons (Fig. 7E), indicating that compact cortical populations contain sufficient information to engage the learned controller and generate coordinated, high-dimensional hand movements.

Together, these findings demonstrate that cortical population activity can directly control an embodied neural network policy, without access to hand-tuned trajectories. By driving the controller through its learned latent representations, the brain-policy interface exploits shared structure between biological and artificial sensorimotor systems, revealing how population dynamics in S1 and M1 can be functionally mapped onto closed-loop motor control.

## Discussion

This study introduces a stimulus-computable, closed-loop framework that links biomechanics, behavioral alignment, and cortical activity during dexterous pre-shaping. Neural network policies trained by imitation learning from observations reproduce naturalistic grasp trajectories and develop internal states that predict both trial-to-trial variability and condition-averaged responses in primary somatosensory (S1) and motor (M1) cortex. Together with the comparative analyses across control modalities and architectures, these results support a view of cortical control as a recurrent, muscle-centric feedback system that integrates proprioceptive and goal signals to achieve robust sensorimotor control. While this view is consistent with the literature (53), we do arrive at it with a monolithic, comprehensive approach.

### Interrogating sensorimotor representation with imitation learning

Our framework advances task-optimized neural modeling (20, 27, 29, 56, 57) by embedding learning within a musculoskeletal, closed-loop system. Unlike prior models that rely on simplified kinematics or direct neural supervision, imitation learning from observations allows controllers to be shaped by realistic physical and feedback constraints that are relevant to sensorimotor computation. Reinforcement learning was essential not as an engineering choice but as a means to discover muscle activation patterns consistent with biomechanical feasibility and behavioral goals, yielding physiologically plausible solutions under non-differentiable dynamics.

The resulting controllers produce smooth, energy-efficient movements that generalize to new objects, suggesting that task optimization under realistic physical constraints can give rise to cortical-like representations (Fig. 2-3). Conceptually, this approach provides a concrete instantiation linking normative ideas from optimal feedback control with embodied models of sensorimotor computation, providing a *stimulus-computable model* that connects sensory goals, proprioceptive inputs, and motor outputs within a unified dynamical system. By showing that neural alignment can emerge from behavioral-alignment alone, the framework offers a datadriven basis for understanding how sensorimotor representations form through embodied interaction with the environment. In parallel work, we have also shown that imitation learning can also produce complex 3D locomotion controlling 80 muscles that aligns with EMG (36).

### Muscle- versus joint-based representations in cortical control

A longstanding question in motor neuroscience concerns the coordinate system underlying movement representation. Classical control theories, including optimal feedback formulations (16, 19, 48, 49), posit joint- or end-effector-centric encoding consistent with robotic control abstractions. In contrast, physiological studies increasingly highlight mixed or muscle-based representations, where cortical populations encode combinations of proprioceptive and efferent variables reflecting the mechanical state of the limb (21, 27, 29, 58, 59). These hypotheses represent two poles of a continuum, from kinematic abstraction to embodied implementation, rather than mutually exclusive hypotheses.

Our results provide model-based evidence relevant to this debate. Although joint-based controllers achieved higher kinematic accuracy, their internal states aligned less closely with cortical activity than those of muscle-based controllers, which directly integrated muscle length, velocity, and force (Fig. 4). This dissociation between behavioral precision and neural alignment suggests that cortical representations may be more consistent with representations supporting coherent integration of motor output and proprioceptive feedback than with purely kinematic descriptions of movement. Because linear baselines using joint or muscle kinematics performed equivalently, the difference likely arises from the representational geometry shaped by closed-loop muscle-level optimization, rather than from differences in input statistics.

The stronger alignment effect observed in M1 compared to S1 supports a functional gradient along the sensorimotor axis. In particular, M1 activity showed greater correspondence with models encoding muscle-level control variables, whereas S1 exhibited weaker but still significant alignment, consistent with hybrid representations combining afferent and efferent information to estimate limb state (7, 12, 20, 37, 60). This pattern reflects recent evidence that muscle-level representations in M1 and S1 arise from recurrent sensorimotor integration during closed-loop control (21). Together, these results argue that the brain’s control architecture emphasizes embodied feedback integration rather than purely geometric mapping between joints and space.

Experimentally, this finding motivates future studies combining intracortical recordings with direct EMG or tendon-force measurements to determine whether the simulated activations correspond to true physiological synergies.

### Adaptive temporal sensorimotor processing

Our architectural analyses reveal how temporal integration shapes sensorimotor representations. Among tested architectures, recurrent networks, particularly LSTMs, showed the strongest alignment with both behavior and neural activity, outperforming feedforward, convolutional, graph-based, and attention-based models (Fig. 5). This advantage underscores the importance of maintaining and updating internal states over short time horizons to integrate proprioceptive feedback and efferent history during dexterous control.

These findings parallel prior work showing that recurrent neural networks trained for reaching or arm control develop low-dimensional, rotational dynamics resembling those observed in motor cortex (27–29, 31, 54). Such dynamics are thought to implement internal feedback control, combining sensory evidence and predictive estimates to generate coordinated trajectories (17, 18). Extending this principle to dexterous grasping, our results indicate that recurrence, rather than instantaneous mapping, captures the temporal dependencies required for fine coordination across many muscles. While static encoders such as MLPs captured steady-state mappings between proprioception and motor output, only recurrent architectures achieved higher explained variance when predicting time-resolved neural activity.

In contrast, GNN and Transformer architectures, though powerful for modeling structured and attention-based relationships (47), may not readily capture the smooth, time-evolving feedback loops characteristic of sensorimotor control. Their emphasis on relational or context-dependent weighting favors flexible feature selection over the continuous temporal integration that defines cortical dynamics. Thus, their weaker alignment suggests that, in this task regime, architectures with recurrent internal state provide a more effective inductive bias for predicting sensorimotor population activity than purely attentional or graph-based aggregation.

### Integration of sensory and motor signals across cortical circuits

Classical frameworks often depict grasping as a serial cascade, from visual and parietal processing to premotor and motor execution (11, 13). Yet growing evidence points to a distributed, bidirectional organization in which sensory and motor cortices interact dynamically (14, 20, 61–64). Our results support this integrative view. Within the controller, the convergence layer where proprioceptive, efferent, and goal streams merge yielded the highest neural predictability in both S1 and M1 (Fig. 6), consistent with shared sensorimotor representations arising from feedback integration.

This convergence is consistent known anatomical motifs linking sensory and motor areas through thalamocortical and cortico-cortical pathways (7, 20). Although S1 and M1 differ in their canonical roles, sensory processing and motor command, respectively, our results indicate that activity in both areas is best explained by representations in which proprioceptive, efferent, and goal-related signals are integrated rather than segregated. Rather than implying a loss of functional specialization, the strong alignment observed at the convergence layer suggests that both S1 and M1 participate in a shared sensorimotor control state that combines reafferent and predictive information to guide ongoing behavior (21, 60). Together, these findings support a unified framework in which dexterous manipulation arises from recurrent, feedback-driven integration across sensorimotor circuits, while preserving distinct sensory and motor emphases across cortical areas.

### Policy and value representations in cortical computation

The distinction between policy and value networks offers a computational lens on how cortical circuits balance execution and evaluation. In reinforcement learning, the *policy* generates motor commands, whereas the *value* estimates expected future reward from the current state, an internal measure of action quality. This separation has been used as a conceptual abstraction of cortico-basal ganglia organization, where cortical regions propose candidate actions and subcortical loops evaluate predicted outcomes (4, 42, 43, 65).

Across monkeys, neural activity in both S1 and M1 aligned more strongly with the value network than with the policy network under natural closed-loop conditions. This pattern indicates that cortical population activity during grasping is better captured by representations that integrate proprioceptive feedback and task context into a predictive estimate of state than by signals corresponding directly to instantaneous motor commands. Such state-evaluative representations are consistent with internal-model and predictive-coding accounts of sensorimotor cortex, in which ongoing activity reflects expected task progression and error-related information (66, 67).

Importantly, when sensory feedback to the policy network was clamped to the experimentally measured kinematics, eliminating compounding imitation error, alignment between the policy network and cortical activity improved substantially, reaching levels comparable to those of the value network. This result indicates that the weaker alignment of the policy under natural feedback does not reflect an intrinsic mismatch between policy representations and cortical dynamics, but rather the sensitivity of action-generating networks to small deviations in closed-loop imitation. Together, these findings suggest that cortical population activity during grasping reflects predictive, state-dependent representations that are compatible with both action generation and evaluative processes, without implying a strict separation of policy and value computations within cortex.

Similar actor-critic architectures have been used as conceptual models of motor sequence control in songbird basal ganglia-thalamocortical loops (68), and dopaminergic projections to motor and somatosensory cortex (43) provide a potential substrate for evaluative modulation of cortical activity. Together, our results support the view that sensorimotor cortical dynamics during dexterous control are best explained by integrated representations that combine prediction, evaluation, and action-related information within a shared recurrent framework.

### Neural decoding and brain-controlled policy

By replacing the reference input with a brain-derived signal, we show that population activity in sensorimotor cortex can steer the controller through its goal pathway. Two interfaces were compared: a classical joint-angle decoder mapping neural activity to kinematics (69–71) and a latent-trajectory decoder mapping neural activity to the policy’s internal trajectory embedding. Both interfaces produced coherent, object-specific grasping behavior above a fixed-posture baseline. Critically, decoding into the learned latent space matched the accuracy of direct kinematic decoding on held-out objects while exhibiting substantially greater tolerance to injected noise. Performance was higher when decoding from M1 than from S1, yet both areas supported effective control, and reliable operation was achieved using activity from only tens of neurons, reflecting the compactness of the learned control representation.

These findings refine a long-standing BMI assumption that cortical activity must be decoded into explicit kinematic variables. Interfacing with a muscle-centric, feedback-aware latent space, which already integrates goals with proprioceptive context, yields an interface that is both accurate and robust. The robustness advantage under Poisson-like perturbations suggests that the policy’s latent embedding acts as a stabilizing control representation that attenuates decoding noise, consistent with evidence that postural or goal-related variables can provide more stable control signals than instantaneous velocity (72). That S1 alone can sustain above-baseline control, albeit weaker than M1, supports a shared sensorimotor code in which afferent state estimates and efferent goals are jointly represented (20). Conceptually, aligning neural activity with learned latent representations illustrates a promising direction for manifold-aligned BMIs that emphasize stability and task relevance over direct kinematic mapping (73).

### Limitations and future directions

Although the present framework provides a biologically grounded model of sensorimotor computation, several limitations remain. The musculoskeletal model employed here, though dynamically validated and widely used for reinforcement learning (74), was derived from the human hand. Following the approach of Goodman et al. (12), we mapped primate joint angles onto this model to preserve dynamical realism and generate meaningful muscle activations. While this approximation captures the qualitative structure of primate grasping, it cannot fully reproduce species-specific anatomy or muscle-tendon routing. Developing primate-specific, GPU-accelerated musculoskeletal models incorporating experimentally measured parameters would increase biological fidelity and enable large-scale training of embodied controllers at neural resolution.

Our analysis also focused on the pre-contact phase of grasping, omitting tactile feedback, which is known to modulate neural responses during object interaction and contact stabilization. Extending the framework to include contact dynamics and tactile sensing would enable direct investigation of somatosensory prediction errors and feedback-driven corrections (21, 53, 60), bridging prehension and manipulation within a single computational model. Likewise, future architectures could introduce hierarchical recurrent organization, reflecting the nested feedback loops observed across spinal, thalamocortical, and parietofrontal circuits. Such models may help clarify how rapid proprioceptive feedback interacts with slower, goal-dependent control processes to maintain stable, adaptive movement.

Finally, our neural analyses focused on single-unit encoding and inferred muscle activations from kinematics rather than physiological recordings. Simultaneous intracortical and EMG measurements during grasp would provide a decisive test of whether the simulated muscle activations correspond to true physiological synergies, and whether the geometry of network latent spaces aligns with population-level neural manifolds (54, 55). These multimodal extensions would close the loop between simulation and experiment, refining both the biological interpretation of our results and the mechanistic grounding of model-based neuroscience.

## Methods

### Behavioral and neural experimental data

We used single-cell neural activity recordings obtained from four primates using Utah arrays by Goodman et al. (12).

#### Behavioral data during grasping movements

Primates are trained to perform a grasping task that required manipulating hand posture to grasp 35 different objects of varying size, shape and orientation. The same object but with a different orientation was considered as a “different” object since it requires a different grasping strategy. Each object was presented eight to eleven times in a given session resulting in approximately 300 trials per session.

At the beginning of the trials, objects are magnetically attached to a robot arm that keep them “out of reach”. Trials are initiated only when primates keep their arms still in the chair’s armrests, monitored by photosensors. After a 1-3 s delay, randomly drawn on a trial-by-trial basis, the robot moves the objects toward the primate hand for grasping. Two main events were identified based on the kinematics: start of movement, the time at which the hand began to move about the wrist joint; and grasp, when object contact was finally established. The interval of interest for both the behavioral and neural data spanned between 750 ms before the start of the movement to the onset of the movement (hold period) and from the movement onset to 10 ms prior the contact with the object (grasp period). More details about the behavioral experiment can be found in Goodman et al. (12).

#### Extracellular electrophysiology recordings in sensorimotor areas

Neural activity was recorded from the primary somatosensory cortex (S1) and primary motor cortex (M1) in non-human primates (NHPs) during hand pre-shaping movements (12). Recordings were conducted at a sampling rate of 100 Hz. All NHPs used their left hand during the task, with electrodes implanted in the right hemisphere. An exception was NHP J, which also had an additional implant in the contralateral hemisphere (defined as NHP JL). Spike trains were processed by convolving the raw neural activity with a Gaussian kernel (40 ms window) to estimate firing rates for each neuron. Trial durations varied, with an average length of 140 ms. Neurons with low activity (average firing rates below 0.01 Hz) were excluded from the analysis to ensure robust neural representation.

#### Inferring joint angles from the grasping task

During the behavioral experiments, 31 keypoints from the forearm to the hand were optically tracked using the Vicon system. The keypoints were used to extract the corresponding joint kinematics using a musculoskeletal model of a human arm (75) implemented in Opensim (76). The model includes 39 Hill-type muscle-tendon actuators crossing the forearm, wrist and the hand. In Goodman et al. (12), the musculoskeletal model was scaled to match the size of each primate and the joint kinematics was computed by performing inverse kinematics from the keypoints. The computed joint angles were subsequently translated into a parallel musculoskeletal model implemented in Myosuite (74) that operates within the MuJoCo simulation environment (39). This transition was necessary to facilitate downstream experiments with artificial agents trained with deep reinforcement learning to control the musculoskeletal system. MuJoCo, compared to OpenSim, offers a computationally efficient simulation platform with speeds up to three orders of magnitude faster (74). The musculoskeletal model of the hand specifically represents the right hand. It is worth noting that the left and right hands are symmetric in their movement patterns. Therefore, projecting joint angles from one hand to the other (adjusting for the appropriate angular sign changes) results in equivalent motions. Table 2 lists all muscle-tendon actuators and joint degrees of freedom included in the Mujoco musculoskeletal hand model.

**Table 2.** Muscles and joints of the musculoskeletal hand model. Each muscle and joint is indexed according to its order in the MuJoCo model (0-based). Group labels indicate anatomical or functional classification. *Abbreviations:* CMC = carpometacarpal; MCP/MP = metacarpophalangeal; IP = interphalangeal; PIP/DIP = proximal/distal IP; D2–D5 = digits (index to little).

| Muscles (39) |  |  | Joints (23) |  |  |  |
| --- | --- | --- | --- | --- | --- | --- |
| Idx | Name | Group | Idx | Name | Articulation | Group |
| 0 | ECRL | Wrist extensor | 0 | pro_sup | Radioulnar | Forearm rotation |
| 1 | ECRB | Wrist extensor | 1 | deviation | Radiocarpal | Wrist deviation |
| 2 | ECU | Wrist extensor | 2 | flexion | Radiocarpal | Wrist flexion/extension |
| 3 | FCR | Wrist flexor | 3 | cmc_abduction | Thumb CMC | Thumb abduction |
| 4 | FCU | Wrist flexor | 4 | cmc_flexion | Thumb CMC | Thumb flexion |
| 5 | PL | Wrist flexor | 5 | mp_flexion | Thumb MCP | Thumb MCP flexion |
| 6 | PT | Pronator | 6 | ip_flexion | Thumb IP | Thumb IP flexion |
| 7 | PQ | Pronator | 7 | mcp2_flexion | Index MCP | Digit 2 MCP flexion |
| 8 | FDS5 | Extrinsic flexor (D5) | 8 | mcp2_abduction | Index MCP | Digit 2 MCP ab/adduction |
| 9 | FDS4 | Extrinsic flexor (D4) | 9 | pm2_flexion | Index PIP | Digit 2 PIP flexion |
| 10 | FDS3 | Extrinsic flexor (D3) | 10 | md2_flexion | Index DIP | Digit 2 DIP flexion |
| 11 | FDS2 | Extrinsic flexor (D2) | 11 | mcp3_flexion | Middle MCP | Digit 3 MCP flexion |
| 12 | FDP5 | Extrinsic flexor (D5) | 12 | mcp3_abduction | Middle MCP | Digit 3 MCP ab/adduction |
| 13 | FDP4 | Extrinsic flexor (D4) | 13 | pm3_flexion | Middle PIP | Digit 3 PIP flexion |
| 14 | FDP3 | Extrinsic flexor (D3) | 14 | md3_flexion | Middle DIP | Digit 3 DIP flexion |
| 15 | FDP2 | Extrinsic flexor (D2) | 15 | mcp4_flexion | Ring MCP | Digit 4 MCP flexion |
| 16 | EDC5 | Extrinsic extensor (D5) | 16 | mcp4_abduction | Ring MCP | Digit 4 MCP ab/adduction |
| 17 | EDC4 | Extrinsic extensor (D4) | 17 | pm4_flexion | Ring PIP | Digit 4 PIP flexion |
| 18 | EDC3 | Extrinsic extensor (D3) | 18 | md4_flexion | Ring DIP | Digit 4 DIP flexion |
| 19 | EDC2 | Extrinsic extensor (D2) | 19 | mcp5_flexion | Little MCP | Digit 5 MCP flexion |
| 20 | EDM | Extrinsic extensor (D5) | 20 | mcp5_abduction | Little MCP | Digit 5 MCP ab/adduction |
| 21 | EIP | Extrinsic extensor (D2) | 21 | pm5_flexion | Little PIP | Digit 5 PIP flexion |
| 22 | EPL | Thumb extensor | 22 | md5_flexion | Little DIP | Digit 5 DIP flexion |
| 23 | EPB | Thumb extensor |  |  |  |  |
| 24 | FPL | Thumb flexor |  |  |  |  |
| 25 | APL | Thumb abductor |  |  |  |  |
| 26 | OP | Thumb intrinsic |  |  |  |  |
| 27 | RI2 | Intrinsic (D2) |  |  |  |  |
| 28 | LU_RB2 | Intrinsic (D2) |  |  |  |  |
| 29 | UI_UB2 | Intrinsic (D2) |  |  |  |  |
| 30 | RI3 | Intrinsic (D3) |  |  |  |  |
| 31 | LU_RB3 | Intrinsic (D3) |  |  |  |  |
| 32 | UI_UB3 | Intrinsic (D3) |  |  |  |  |
| 33 | RI4 | Intrinsic (D4) |  |  |  |  |
| 34 | LU_RB4 | Intrinsic (D4) |  |  |  |  |
| 35 | UI_UB4 | Intrinsic (D4) |  |  |  |  |
| 36 | RI5 | Intrinsic (D5) |  |  |  |  |
| 37 | LU_RB5 | Intrinsic (D5) |  |  |  |  |
| 38 | UI_UB5 | Intrinsic (D5) |  |  |  |  |

#### Inferring muscle kinematics with inverse kinematics

As our objective is to use the behavioral data to test hypotheses about the sensorimotor system. We simulated proprioceptive inputs (muscle length and velocity) by setting the estimated joint kinematics to the hand musculoskeletal model. At each timestep, we extracted the corresponding resting muscle length and subsequently computed the first derivative to obtain the muscle velocity.

#### Inferring muscle kinematics with inverse dynamics

While inverse kinematics provides muscle lengths and velocities, it does not capture the muscle dynamics, i.e., the forces and activations required to produce those kinematics. This distinction is essential because, under the Millard muscle model implemented in MyoSuite, muscle force depends nonlinearly on both muscle length and contraction velocity through the force-length-velocity (FLV) relationship. Thus, the forces that would arise during active movement cannot be inferred directly from IK-based kinematics alone. Recovering these dynamics is critical for evaluating whether cortical representations reflect muscle activation, muscle force, or both, as suggested by prior work in primate motor cortex (12).

To obtain physiologically plausible muscle forces and activations associated with the recorded grasping movements, we performed inverse dynamics using the MyoSuite musculoskeletal model. At each timestep, we set the joint kinematics at time *t*+1 and computed the joint torques required to achieve that state using MuJoCo’s inverse dynamics. We then solved for the muscle forces that best reproduced these torques by minimizing the mean-squared error between the predicted and required joint moments:

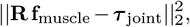

where **R** is the matrix of muscle moment arms (directly available from the MuJoCo model) and **f**_muscle_ contains the unknown muscle forces.

Muscle forces were constrained by the Millard model’s FLV curve. For each muscle, MuJoCo provides the actuator-tendon length

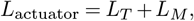

allowing us to compute normalized muscle length,

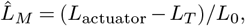

and normalized muscle velocity 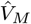. Total muscle force is given by the sum of active and passive components:

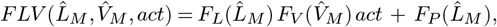

where *act* is the muscle activation state. Activation followed first-order activation dynamics driven by the control signal *ctrl*:

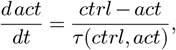

with activation and deactivation time constants governed by the Millard formulation:

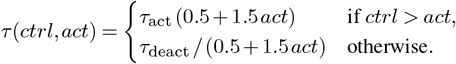

We substituted the FLV-expression for **f**_muscle_ into the optimization problem and solved for the control inputs *ctrl* using quadratic programming, yielding activations and forces consistent with the reference kinematics. Because the musculoskeletal model runs at 500 Hz, joint trajectories (recorded at 100 Hz) were first upsampled to match simulation frequency. Estimated muscle variables were then downsampled back to 100 Hz.

Our implementation builds directly upon the inverse-dynamics procedure provided in the MyoSuite repository (available in the MyoSuite tutorial: https://github.com/MyoHub/myosuite/blob/main/docs/source/tutorials/6_Inverse_Dynamics.ipynb).

### Hierarchical clustering

To compare object-specific representations in both neural activity and joint kinematics, we performed hierarchical clustering analyses based on pairwise dissimilarities between object conditions similar to what was previously done by Schaffelhofer and Scherberger (11). Separate analyses were conducted for neural and kinematic datasets, given their distinct feature spaces across sessions. We focused on the grasp window of the trajectory (from movement onset to the end of the trial). For each object and epoch, neural firing rates and joint angles were averaged across time and repetitions, yielding a single vector per condition. Neural analyses were performed separately for each session and cortical area (S1, M1), while kinematic analyses used concatenated data across sessions.

Joint angles were first z-scored within session to remove calibration and scale differences across days. The condition-mean joint vectors (23 dimensions per condition) were concatenated across sessions and reduced using principal component analysis (PCA) to 10 main components. Pairwise dissimilarities between conditions were computed using the Mahalanobis distance, and hierarchical clustering was performed using average linkage with optimal leaf ordering. The resulting dendrograms reveal similarity structure among object conditions in joint space.

Because the number of neurons differed across sessions, we employed a representational similarity analysis (RSA) approach that avoids explicit neuron alignment. For each session, we computed a condition-by-condition (objects) representational dissimilarity matrix (RDM) using correlation distance between condition-mean firing-rate vectors. Each session-level RDMs were averaged element-wise to obtain a cross-session RDM. Hierarchical clustering was then applied to this averaged RDM using the same average-linkage criterion and optimal leaf ordering. This approach yields a session-agnostic clustering structure that reflects stable representational relationships across recording days.

### Imitation learning task

We designed a trajectory-imitation task in which a policy learns to reproduce joint kinematics observed during primate grasping. Unlike standard imitation learning, where a student policy has access to paired observation-action data (77), here the policy must learn to reproduce behavioral trajectories using only kinematic demonstrations (41). This formulation, *imitation from observations only*, requires the network to infer latent muscle activations that give rise to the observed motion, posing a biologically realistic credit-assignment problem.

The task was designed to generalize beyond individual subjects. At the beginning of each episode, a reference trajectory was uniformly sampled from the full behavioral dataset encompassing multiple primates, objects, and trials. The initial state of the musculoskeletal model was randomized along the selected trajectory (reference-state initialization, RSI), exposing the policy to diverse initial conditions and preventing overfitting to specific kinematic contexts (25, 35). Each episode lasted 1500 ms; if the reference trajectory was shorter, the policy was required to maintain the final pose until the end of the episode. Episodes terminated early if the Euclidean distance between predicted and reference joint positions exceeded 100 mm, preventing unstable exploration.

To emulate a full sensorimotor loop, the MyoSuite environment was modified to use biologically grounded proprioceptive inputs and muscle activations as control signals. Rather than receiving joint angles and velocities, the policy observed surrogate signals analogous to those available through muscle spindles and Golgi tendon organs. The observation vector at each timestep consisted of:

- **Proprioceptive inputs:** muscle lengths (*L*), velocities 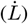, and forces (*F* ) for all 39 muscle-tendon actuators;
- **Previous activation:** muscle activations from the preceding timestep (*a*_*t*−1_), providing an efference copy for dynamic feedback;
- **Reference horizon:** a short look-ahead trajectory of joint poses over a fixed horizon *H*, denoted *obs*_ref_[*t*] = *θ*_ref_[*t* + 1 : *t* + 1 + *H*].

For the main analyses, the reference horizon was set to *H* = 10 control steps (100 ms), allowing the policy to anticipate near-future movements while maintaining computational efficiency.

The policy’s objective was to maximize the expected cumulative reward, balancing trajectory accuracy and energy efficiency to generate smooth, naturalistic movements. The dense step-wise reward was defined as:

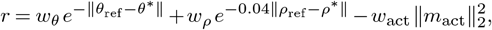

where *θ*_ref_ and *ρ*_ref_ are reference joint angles and keypoints, *θ*^∗^ and *ρ*^∗^ are the corresponding predicted values, and *m*_act_ are muscle activations. The weights were set to *w*_*θ*_ = 1, *w*_*ρ*_ = 1, and *w*_act_ = 10^−3^, prioritizing kinematic fidelity while penalizing excessive activation. This formulation encourages accurate, smooth, and energy-efficient control, mirroring the precision and effort minimization observed in biological grasping.

### Policy training and evaluation

#### Reinforcement learning framework

The policy was trained to imitate experimental kinematic data by framing the problem as a continuous control task within a reinforcement learning (RL) paradigm. We used Proximal Policy Optimization (PPO) (78), an on-policy algorithm well suited for continuous action spaces and stable gradient updates. Training and rollout collection were implemented using the Stable Baselines 3 library (79), with custom environment wrappers to support muscle-level control and trajectory-based imitation (74).

The policy followed an actor-critic architecture, where the actor predicted muscle activations at each timestep and the critic estimated the expected reward to guide policy updates. To enhance exploration in the overactuated musculoskeletal system, we employed Lattice exploration (24), which exploits correlations among actuators to sample structured activation patterns. Both actor and critic networks were trained independently but shared the same architecture and optimization hyperparameters.

Optimization was performed using the Adam optimizer with a learning rate of 2.55 *×* 10^−5^. Training ran for 5 *×* 10^7^ environment steps, corresponding to approximately 6 *×* 10^3^ policy updates. Rollouts were collected in segments of 128 steps, and minibatches of 256 samples were used per gradient update. PPO-specific parameters were set following best practices for continuous control: clipping parameter 0.3, discount factor *γ* = 0.99, generalized advantage estimation (GAE) *λ* = 0.9, gradient clipping at 0.7, and entropy regularization coefficient 3.6 *×* 10^−6^. Each update comprised 10 optimization epochs, and the value-function loss coefficient was set to 0.836. Observation normalization was applied to all input streams at every update step.

All experiments used 64 parallel environment instances for on-policy rollouts and were trained on NVIDIA A100 GPUs using PyTorch 2.2 and MuJoCo 3.1.1. The full hyperparameter configuration is provided in Table 3.

**Table 3.** Training hyperparameters for the PPO-based policy.

| Parameter | Value |
| --- | --- |
| Algorithm | Proximal Policy Optimization (PPO) |
| Implementation | Stable Baselines 3 (PyTorch 2.2) |
| Environment instances | 64 parallel rollouts |
| Total timesteps | $5 \times 10^7$ |
| Rollout length (per env) | 128 steps |
| Batch size | 256 samples |
| Optimization epochs per update | 10 |
| Learning rate | $2.55 \times 10^{-5}$ |
| Optimizer | Adam ($\beta_1 = 0.9$, $\beta_2 = 0.999$) |
| Discount factor ($\gamma$) | 0.99 |
| GAE parameter ($\lambda$) | 0.9 |
| Clip range | 0.3 |
| Entropy coefficient ($\beta$) | $3.6 \times 10^{-6}$ |
| Value-loss coefficient ($c_v$) | 0.836 |
| Gradient clipping (norm) | 0.7 |
| Observation normalization | Enabled |
| Lattice exploration | Enabled (24) |
| Hardware | NVIDIA A100 (80 GB) GPU |

#### Policy evaluation

The performance of each trained policy was assessed by comparing predicted and reference hand kinematics across all primates. Specifically, we computed the mean Euclidean distance between predicted and reference keypoint positions in Cartesian space. The musculoskeletal model includes 36 keypoints spanning the forearm and hand, from which we selected a subset of 23 hand keypoints to focus the evaluation on dexterous control.

For each trial, Euclidean distances were averaged across keypoints and then across time, yielding a single trajectory-level score. The overall performance metric was obtained by averaging these values across all trials and subjects. We additionally report a *relative error* metric obtained by normalizing the Euclidean distance by the pinky fingertip length. This normalization provides a dimensionless measure of tracking accuracy.

#### Policy robustness

To assess robustness and identify the functional contribution of each sensory stream, we introduced controlled perturbations to the inputs during evaluation. Each perturbation type was applied independently to isolate specific effects:

##### (a) Signal-dependent noise

Gaussian noise with a standard deviation of 10% of the signal magnitude was added to test resilience to natural sensory variability.

##### (b) Zeroed input

A given sensory modality was replaced with zeros to assess dependence on that information channel.

##### (c) Constant input

Signals were fixed to their initial trial values, testing adaptability to static yet realistic sensory feedback.

##### (d) Temporal delay

A 60 ms lag was applied to simulate physiological sensorimotor delays and test the importance of real-time feedback.

Policy performance under each perturbation was evaluated using the same Euclidean distance metrics described above. Comparisons across conditions revealed the sensitivity of the learned controller to sensory degradation and latency, providing insight into the relative weighting of proprioceptive, efferent, and goal inputs during control.

To complement these test-time manipulations, we performed ablation experiments during training in which individual sensory modalities were removed entirely from the observation space. The resulting change in final performance quantified each modality’s causal contribution to learning, distinguishing features that support robust adaptation from those that primarily stabilize behavior post-training.

### Neural network architecture

Building on the sensorimotor formulation described above, we implemented a family of actor-critic architectures designed to test competing hypotheses about how proprioceptive, efferent, and goal-related signals are processed and integrated. Each sensory stream was encoded by a dedicated subnetwork, allowing for modular and biologically interpretable representations without shared weights across modalities.

#### Architectural organization

Each model consisted of two functional components: (i) independent stream-specific encoders that processed sensory and motor-related inputs, (ii) a multimodal integration stage (decoder) where latent features were concatenated and produced either muscle activations (policy) or a scalar value estimate (critic). Unless otherwise noted, encoders had 128 hidden units and decoders 256 per layer, with ReLU activations. Weights were initialized with Xavier uniform initialization. These architectural constants were fixed across model variants to isolate the contribution of each computational mechanism.

To investigate distinct hypotheses about spatial and temporal integration, we systematically varied the encoder and decoder modules. All architectures were trained under identical imitation- and reinforcement-learning regimes. Table 4 summarizes the full set of evaluated configurations. The different architectures are:

**Table 4.**
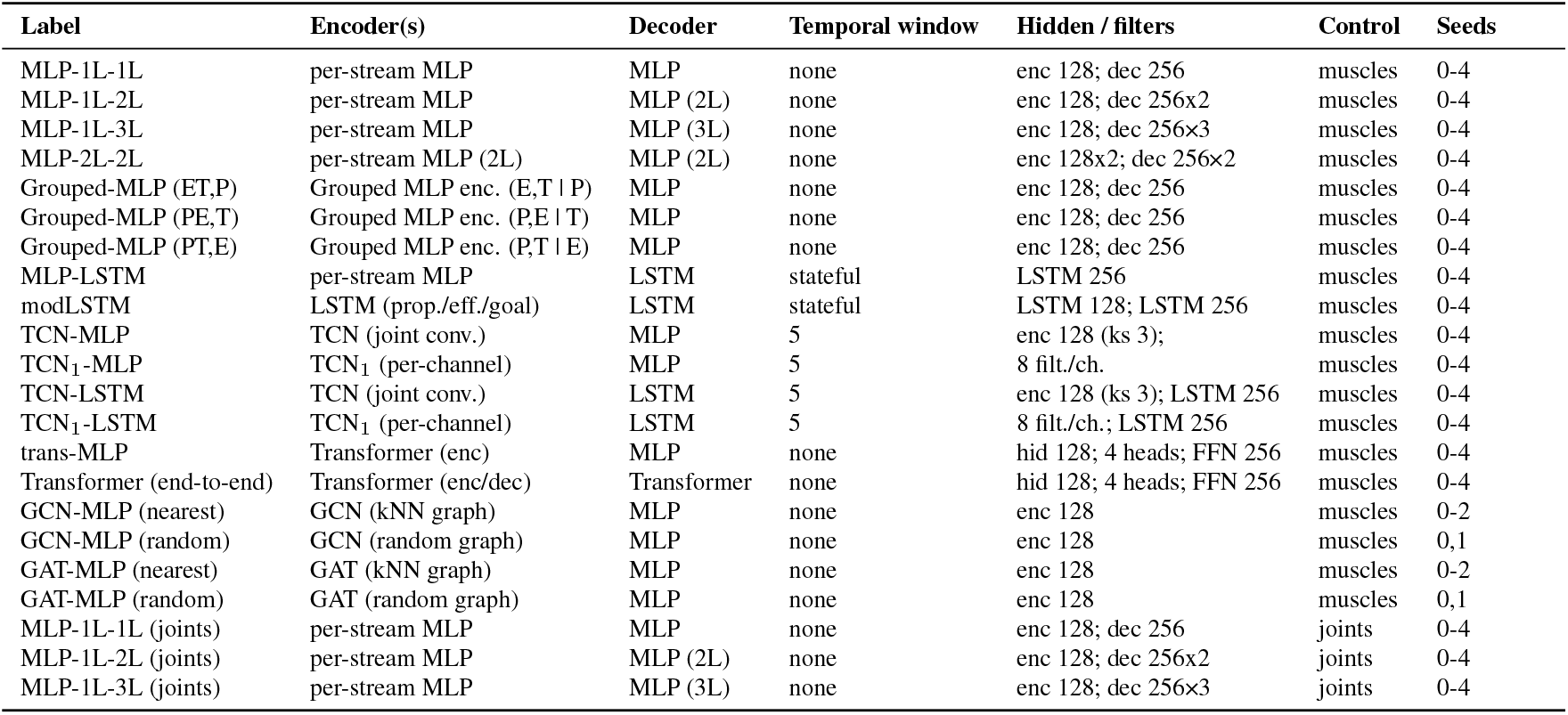
Model inventory and key architectural parameters. Abbrev.: TCN_1_ = per-channel temporal conv.; modLSTM = per-stream LSTM; trans = Transformer encoder; GCN/GAT = graph encoders; grouped MLP = hierarchical MLP in which selected streams are first encoded jointly, then integrated at a higher level. Integration notation: (E,T | P) = integrate motor efference and trajectory encoder before proprioception; (P,E | T) = integrate proprioception and efference before the trajectory encoder; (P,T | E) = integrate proprioception and the trajectory encoder before efference. Unless noted: encoder hidden = 128, decoder hidden = 256 ( *× L*), control step = 10 ms, horizon *H* = 10, seeds = 5, and kernel size (ks) = 3 for TCN-based models.

#### MLP networks

Baseline fully connected models in which each sensory stream was encoded by a one-layer MLP (or two). The decoder contained one to three layers. This configuration implements global, all-to-all coupling between muscles, analogous to distributed population activity in motor cortex (10, 12).

#### Recurrent networks (LSTM, modLSTM)

To model adaptive temporal integration, the decoder was replaced by a single-layer LSTM. The *modLSTM* variant extended this design by assigning an independent LSTM encoder to each sensory stream (proprioceptive, efferent, goal), whose hidden states were concatenated and passed to a recurrent decoder LSTM. This hierarchical recurrence tests whether maintaining internal states over time facilitates temporal credit assignment and predictive motor behavior (27, 29).

#### Graph neural networks (GCN, GAT)

These models tested whether imposing an explicit muscle–muscle topology influences representation learning. Each muscle was represented as a node in a graph. In the k-nearest configuration (*k* = 1), every muscle was connected to its immediate neighbor in index order, forming a simple chain graph. In the random configuration, each muscle was connected to a randomly chosen partner, preserving one edge per node but disrupting anatomical structure. The *GCN* variant applied a degree-normalized mean aggregation over neighboring muscles, whereas the *GAT* variant replaced this fixed averaging with learned attention weights that modulated each neighbor’s contribution. Both graph encoders were followed by an MLP decoder. Random-topology controls tested whether performance depended on the structured connectivity of the muscle graph (45).

#### Temporal convolutional networks (TCN, TCN_1_)

To model fixed-window temporal dependencies, each stream included the previous five timesteps (*T* = 5). The *TCN* applied joint 1D convolutions (kernel size 3, stride 1) across all features, while the *TCN*_1_ applied channel-wise convolutions followed by concatenation that was fed into the MLP decoder. These models approximate short-latency feedback dynamics.

#### Transformer networks

Attention-based architectures were implemented to capture dynamic, context-dependent reweighting of proprioceptive and efferent signals. Two configurations were used. In the *Transformer-MLP* model, proprioceptive (length, velocity), force, activation, and goal inputs were first embedded independently. The encoder consisted of a single Transformer block with learned positional encodings representing muscle identity and layer normalization applied before each sub-layer (*ϵ* = 10^−5^). Self-attention was performed across muscles to capture structured intermuscle dependencies, and the resulting embeddings were concatenated with the goal representation and passed through a single-layer MLP to generate muscle activations.

In the *Transformer-only* configuration, both encoder and decoder were Transformer blocks with identical hyperparameters. Self-attention was first applied across muscle-level tokens, followed by cross-attention between activation queries and the proprioceptive-goal context, yielding a fully attention-driven policy capable of integrating sensory and goal information end-to-end.

To ensure compatibility with the Transformer input format, the environment’s observation pipeline was modified to output structured token sequences. Each token represented a muscle-level sensory channel comprising spindle-like inputs (length, velocity), tendon force, and efferent activation, together with task-goal features representing future joint configurations. Learned positional embeddings indexed muscle identity, enabling the Transformer to attend across spatially organized proprioceptive features while maintaining a biologically grounded sensorimotor organization.

#### Hierarchical temporal architectures

Combinations of convolutional encoders (TCN or TCN_1_) and recurrent decoders (LSTM) were included to test whether hierarchical temporal processing, fixed local filtering followed by adaptive memory, improves behavioral performance and neural alignment.

### Vision-conditioned policy model

#### Overview

To test whether the controller could generalize beyond oracle (experimentally derived) trajectories, we trained a diffusion-based generative model capable of reconstructing grasping movements directly from visual object information. This approach allowed us to replace ground-truth kinematics with realistic, vision-conditioned trajectories, providing a fully self-contained framework that links visual perception, trajectory generation, and motor control. We modeled grasping hand motion as a conditional diffusion process over trajectories of joint angles. We adapted the approach of Human Motion Diffusion Model (MDM; (80)): (i) a transformer encoder backbone, (ii) classifier-free guidance (CFG; (81)), and (iii) direct prediction of the clean sample **x**_0_ at each reverse step. Unlike MDM, which conditioned on text, we conditioned on visual object features to generate object-specific grasp trajectories.

Let **x**_0_ ∈ ℝ^*L×D*^ denote a clean trajectory of length *L* with *D* = 23 joint angles per time step. A forward diffusion process added Gaussian noise over *T* = 1000 steps, producing **x**_*t*_ according to a cosine variance schedule. At reverse timestep *t* the network *G*(**x**_*t*_, *t*, **c**) predicted 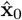, where **c** was the object conditioning vector and **0** denoted the null condition. Classifier-free guidance formed a guided clean prediction

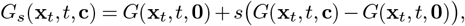

with guidance scale *s* = 3.0 as in MDM.

#### Egocentric object view

Because no calibrated multi-view recordings of the grasped objects were available, we constructed an egocentric visual dataset in Blender (82) to approximate the viewpoint of the macaque during pre-shaping.

Each of the 35 objects used in the behavioral task was manually modeled in Blender based on the reference photographs acquired during experiments. Exact physical dimensions were not available, so object geometries were scaled through qualitative estimation from these images.

To generate visual inputs for the diffusion model, we rendered each object from 100 slightly perturbed camera poses centered around a frontal, near-egocentric viewpoint. The camera distance, azimuth, and elevation were jittered within a small range to mimic the natural variability of head-centered viewpoints while keeping the object approximately centered and upright. This procedure produced a dataset of 3’500 rendered RGB images (100 per object). These renders were paired with the corresponding object class labels from the Goodman et al. (12) dataset, which comprised 7,371 recorded grasping trajectories across 3 monkeys and 21 sessions (approximately 210 trajectories per object).

During training, a matched object render was sampled uniformly at random for each trajectory. Because the true viewpoint of the macaque at grasp onset is unknown, randomizing the render for each training sample reduced spurious correlations between specific images and specific trajectories, encouraging the model to learn object-specific grasp affordances rather than image–trajectory memorization.

#### Representation and conditioning

Each pose (23 joint angles) was linearly projected to a latent embedding of dimension *d* = 512. A sinusoidal positional encoding was added to capture temporal order. The diffusion timestep *t* was embedded via a 2-layer MLP acting on a sinusoidal timestep encoding, producing a *d*-dimensional vector that was added to every token. For visual conditioning, a frozen ResNet-50 backbone (83) processed a rendered object image, whose features were then linearly projected to **c** ∈ ℝ^*d*^. This conditioning vector was broadcast and added to the token embeddings, providing global object context.

#### Architecture

The denoiser *G* was a 6-layer transformer encoder (84) (8 attention heads, hidden size 1024, GELU activation, dropout 0.1). The output embedding at each time step was linearly mapped back to *D* joint angles, yielding the clean trajectory prediction 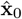. Padding masks were propagated through attention layers to ignore padded frames.

#### Diffusion process

We used a cosine variance schedule that produced monotonically increasing *β*_*t*_ ∈ (0, 1), with *α*_*t*_ = 1 − *β*_*t*_ and cumulative products 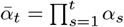. Uniformly sampling *t* ∼ 𝒰*{*1,…, *T}*, the noisy sample was:

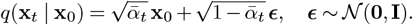

Following Tevet et al. (80), in the reverse step the model directly predicted the clean trajectory **x**_0_ rather than the noise. The posterior *q*(**x**_*t*−1_ | **x**_*t*_, **x**_0_) was Gaussian with mean

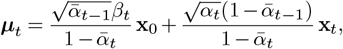

and variance proportional to *β*_*t*_ (standard DDPM formulation; (85)). At inference we ran the reverse diffusion for *T* = 1000 steps. For each *t* we formed both conditional and unconditional predictions, generated the guided clean prediction *G*_*s*_(**x**_*t*_, *t*, **c**) with *s* = 3.0, computed ***µ***_*t*_, sampled and proceeded to *t*− 1.

#### Policy training with diffusion-generated trajectories

Once trained, it generated novel, object-specific grasp trajectories from single egocentric images. These diffusion-generated trajectories were subsequently used to condition the policy networks, allowing us to evaluate controller performance when trained on visually inferred rather than oracle trajectories (see “Policy training and evaluation”).

### Comparison between muscle- and joint-based controllers

To examine how different levels of motor abstraction influence behavioral performance and neural alignment, we trained two classes of policies that differed only in their control and observation modalities: *muscle-based controllers*, which operated directly on muscle activations, and *joint-based controllers*, which acted at the level of kinematic degrees of freedom. All other components of the training pipeline, including the architecture, learning algorithm, reward formulation, and data sampling, were identical across models.

For the joint-based controllers, the musculoskeletal XML model was modified by removing the muscle actuators and by adding torque actuators to each of the 23 kinematic joints, with control ranges set to [ −1, 1] and unit gear ratios. The proprioceptive observation space was defined in joint coordinates rather than muscle space, comprising joint angles, angular velocities, and net torques for all 23 degrees of freedom. In addition, the policy received the reference trajectory horizon and previous net torques history. The policy output consisted of 23 continuous torque commands, which were applied directly to the corresponding joint actuators at each control step.

The reward function, PPO training parameters, and rollout configuration were identical to those used for muscle-based policies (see “Policy training and evaluation”). Network architectures were adjusted only to match the dimensionality of the input and output layers. For the joint-based condition, we trained a total of 15 MLP models comprising one encoder layer and between one and three decoder layers, each evaluated over five random seeds.

Neural alignment was quantified using the same encoding-analysis pipeline (see “Neural predictions”), ensuring direct comparability between joint-based and muscle-based controllers. This approach allowed us to isolate the representational consequences of controlling movement at different hierarchical levels while keeping all optimization and training conditions constant.

### Neural predictions

#### Neural network activations extraction

To analyze the internal representations of the trained controller, we employed two complementary approaches that differed in the source of sensory inputs: (a) Dynamic representations. Sensory inputs arose naturally from the policy’s closed-loop interaction with the simulated environment. These inputs reflected the proprioceptive feedback generated by the network’s control actions, evolving dynamically as the agent interacted with its musculoskeletal model. This approach mirrors the natural emergence of proprioceptive signals during movement. (b) Clamped representations. Sensory inputs were clamped to values derived from inverse kinematics (IK) computed from the experimental behavioral data. This ensured that the network received the same sensory trajectories observed in the real experiments, aligning our analysis with standard approaches in the neural modeling literature.

For both approaches, activations from each network layer were extracted over time and across all trials. Principal component analysis (PCA) was applied to reduce the dimensionality of these activations to the first 78 principal components (PCs), matching the number of regressors used in the baseline linear models (see below). This dimensionality reduction retained the majority of variance in each layer’s activity while ensuring consistent input dimensionality across models. The resulting PCs were used as predictors to model single-neuron firing rates recorded in S1 and M1.

#### Single-neuron prediction models

Single-neuron activity was modeled using ridge regression from the 78 PCs of each layer’s activations. Unlike traditional time-varying encoding models, we trained a single linear regression per neuron with fixed coefficients across time. In other words, the mapping between network activations and neural firing rates was static within each neuron, under the assumption that temporal dynamics are already captured in the network’s internal states. This fixed mapping enables generalization across entire test trials without requiring refitting at each timestep.

For each recording session, trials were split by object identity into 80% training and 20% testing sets, based on complete trials rather than within-trial samples. This trial-based splitting represents a more challenging evaluation and it better reflects the biological constraints of continuous motor behaviors. Model generalization was evaluated separately for (i) IID test trials from the same objects as the training set and (ii) OOD test trials containing previously unseen objects. For each neuron, we trained independent ridge regression models for every network layer. The regularization parameter (*α*) was optimized via 3-fold cross-validation on the training data over 15 logarithmically spaced values between 10^−2^ and 10^5^, and the final model was refit using the full training set.

#### Condition-averaged and single-trial neural predictions

To assess how well network states captured both trial-specific fluctuations and condition-averaged tuning, we evaluated two complementary prediction regimes: (a) Single-trial predictions. Ridge models were used to predict neural activity at every timestep of each individual trial in the test set. The resulting predicted firing rates were compared to the recorded single-trial activity, and explained variance (EV) was computed independently for each neuron. This approach assesses whether the model can reproduce moment-to-moment variability within individual grasping movements. (b) Condition-averaged predictions. Each trial is composed of a 750 ms hold phase and, after movement onset, a variable-length grasp phase. To ensure comparable temporal alignment across repetitions of the same object, The grasp phase of each trial was linearly interpolated to match the maximum duration observed across all trials of that object, resulting in condition-averaged neural and predicted activity with consistent temporal length. Neural and predicted signals were then averaged across trials for each condition, and explained variance was computed between the condition-averaged predicted and recorded responses. This approach captures the model’s ability to reproduce stable condition-specific tuning while preserving temporal structure within the movement sequence.

#### Baseline encoding models

As a comparison, we trained standard linear encoding models predicting single-neuron firing rates from kinematic variables (muscle- or joint-based, depending on the control architecture). These models used the same cross-validation procedure and hyperparameter optimization as the network-based regressions, providing direct baselines for evaluating task-driven representational alignment.

#### Neural prediction metrics

Prediction accuracy was quantified using the fraction of explained variance (EV):

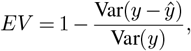

where *y* and *ŷ* denote the observed and predicted firing rates, respectively. For each neuron, we computed EV values for all layers and test conditions (IID and OOD). To summarize model performance, we used the average best layer EV across neurons:

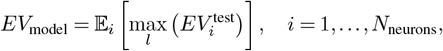

where *l* indexes the network layer yielding the highest predictability for each neuron. This approach identifies the most brain-like representational stage for each neuron without assuming a fixed correspondence between layers and cortical areas. In addition, we evaluated layer-specific EV distributions within S1 and M1 to examine the hierarchical correspondence between model depth and cortical organization.

### Brain-controlled policy

The trained policy was designed as a feedback controller conditioned on a short horizon of future reference trajectories. To test whether neural activity could directly drive this controller, we implemented two linear decoders that mapped population activity from sensorimotor cortex (S1 or M1) onto task-relevant control features used by the policy. (1) *Joint-angle decoder*. This decoder mapped neural population activity to joint angles, following conventional brain-machine interface (BMI) approaches where cortical firing rates are translated into kinematic variables. (2) *Latent-representation decoder*. This decoder mapped neural activity to the principal components (PCs) of the trajectory encoder’s latent representation within the trained policy. Because these latent embeddings exhibited strong alignment with cortical activity, decoding into this internal representational space was hypothesized to yield more robust and behaviorally meaningful control signals. Both decoders were implemented using ridge regression with the same training and test splits as in the neural predictivity analysis. Neural firing rates were used as regressors, and the target variables were either joint angles or latent trajectory embeddings. The linear regression defined a fixed, time-invariant mapping between neural activity and control features allowing the models to generalize across trials.

Each trained decoder was embedded within the imitation-trained policy, replacing the reference trajectory input with real or simulated neural activity. The resulting *brain-controlled policy* therefore used decoded cortical signals, rather than kinematic references, to generate muscle activations and interact dynamically with the musculoskeletal environment. We assessed control performance by computing the egocentric distance between the policy-generated trajectories and the corresponding behavioral trajectories. To evaluate the stability of the neural interface, we tested the robustness of both decoders to noise in the input neural activity. Specifically, we injected Poisson-scaled noise proportional to the firing rate magnitude and measured its effect on closed-loop performance. This analysis quantified how decoding accuracy, and consequently behavioral control, degraded with increasing noise levels. Decoding into the policy’s latent trajectory embedding yielded substantially greater noise tolerance than direct joint-angle decoding, indicating that the learned latent representations capture more stable and noise-resistant control features. Together, these analyses demonstrate that cortical population activity can be transformed into effective control commands for a learned motor policy. Decoding through the model’s latent representation provides a robust and biologically meaningful pathway for brain-driven control, linking cortical population dynamics with closed-loop sensorimotor computation.

## Acknowledgments

We are indebted to Sliman Bensmaia for sharing the data from the Goodman et al. study. We thank members of the Mathis Group for feedback during this project.

## Author contributions

A.M., A.M.V. and M.W.M conceived the project. A.M.V. implemented the models and carried out all analyses with feedback from all authors. A.M.V., and

A.M. wrote the manuscript with input from all authors. A.M. supervised the project and acquired funding.

---

**Fig. S1.**
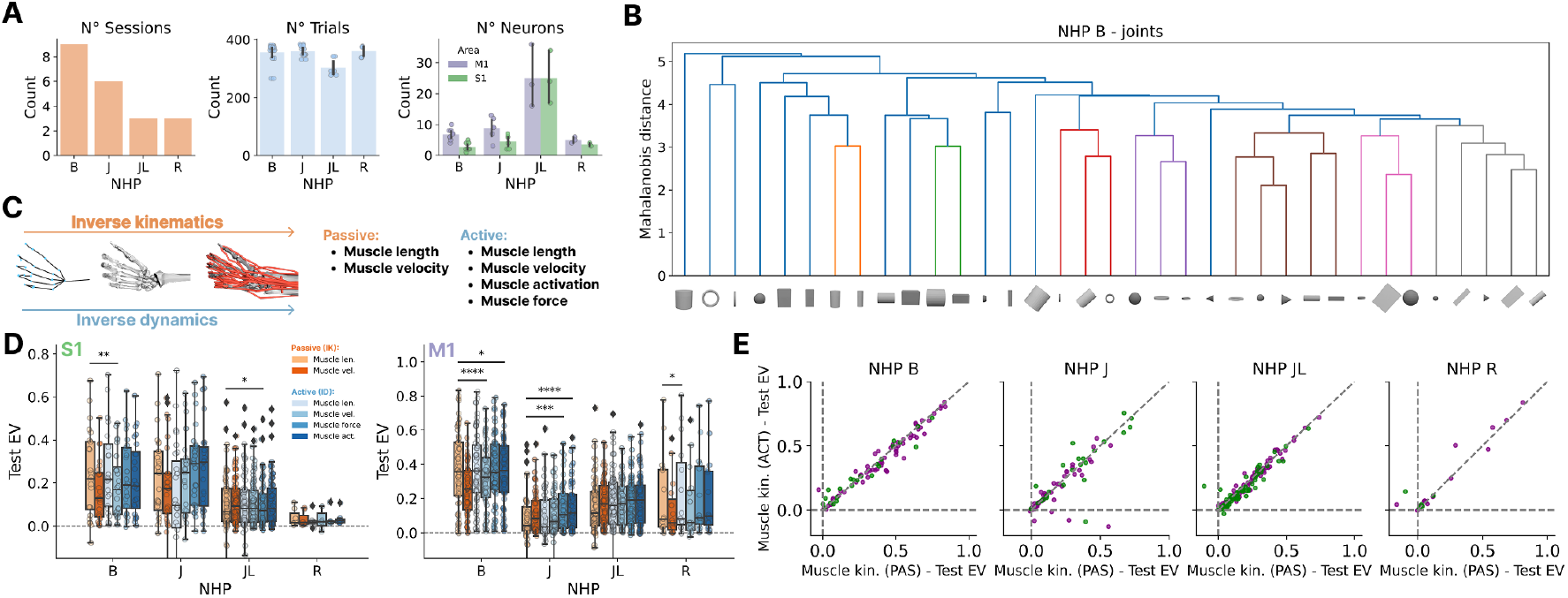
Overview of neural recordings and muscle-based encoding models. **A**. Summary of the neural dataset. Number of recording sessions per non-human primate (NHP; left), number of grasping trials per session (center; each dot represents a session), and number of simultaneously recorded neurons in somatosensory (S1, green) and motor (M1, purple) cortex for each NHP (right). **B**. Hierarchical clustering of object-specific joint configurations for one NHP (B), showing structured relationships among grasp postures. **C**. Schematic of the inverse-kinematics (IK, “passive”) and inverse-dynamics (ID, “active”) pipelines used to estimate muscle kinematics and forces from joint trajectories. IK-derived variables include muscle length and velocity, while ID additionally yields muscle activation and force. **D**. Comparison of neural predictions using individual active and passive muscle variables on out-of-distribution (OOD) test objects. Muscle activations and forces achieve similar EV to muscle length and velocity for both cortical areas, indicating that active and passive muscle features contribute comparably to explaining neural activity. **E**. Distributions of single-neuron prediction accuracy using muscle kinematics as regressors, shown separately for S1 (left) and M1 (right). M1 neurons exhibit consistently higher predictability than S1.

**Fig. S2.**
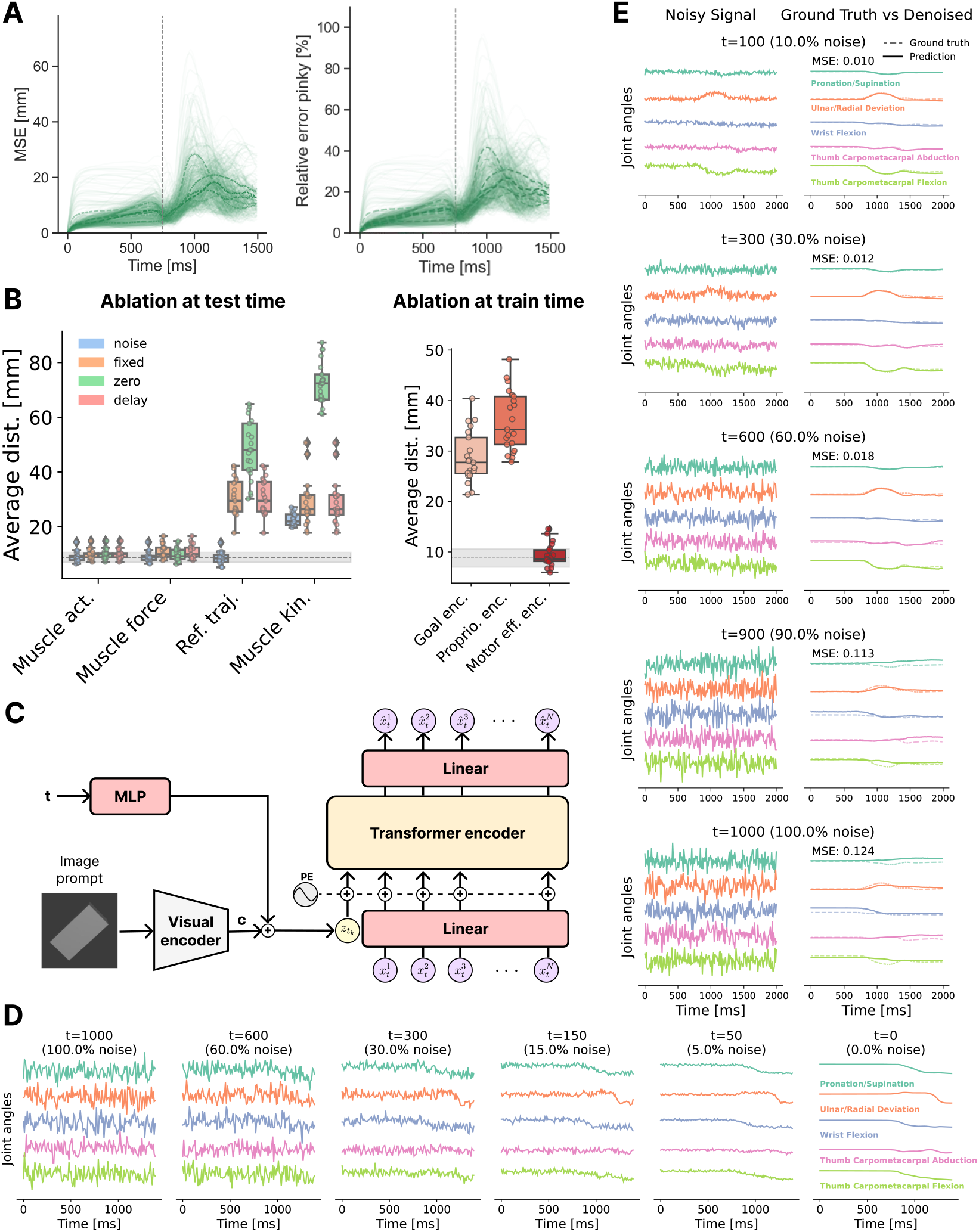
**A**. Egocentric distance over time for all out-of-distribution (OOD) trajectories within a session, averaged across keypoints (left). Peak errors occur around movement onset. Right: egocentric distance normalized by the pinky fingertip length, providing a scale-invariant measure of error. Both measures show consistent temporal profiles across objects. **B**. Left: Distribution of task performance for the 1-layer controller under test-time perturbations (*noise, fixed, zero*, and *delay* ), aggregated across sessions and non-human primates (NHPs). The gray shaded area indicates baseline performance without perturbations. Right: Performance when specific input streams were removed during training (*reference trajectory, proprioception*, or *previous activations*). Both analyses show that proprioceptive and goal-related inputs are essential for accurate pre-shaping. **C**. Architecture of the vision-conditioned diffusion model. The model adapts the Human Motion Diffusion Model (MDM) (80) for our dataset by replacing text-conditioning with visual conditioning. Visual embeddings are extracted from a single egocentric image using a ResNet50 encoder, linearly projected to the transformer token dimension, and combined with standard MDM time and positional encoders. The network directly regresses denoised trajectories at each diffusion step. **D**. Intermediate outputs from the diffusion model across timesteps (*t* = 1000 to *t* = 0), showing progressive refinement of initially noisy trajectories into smooth, coherent pre-shaping motions. **E**. Evaluation of the trained diffusion model on OOD test trajectories, illustrating accurate denoising and reconstruction of behavioral kinematics across diverse grasp types.

**Fig. S3.**
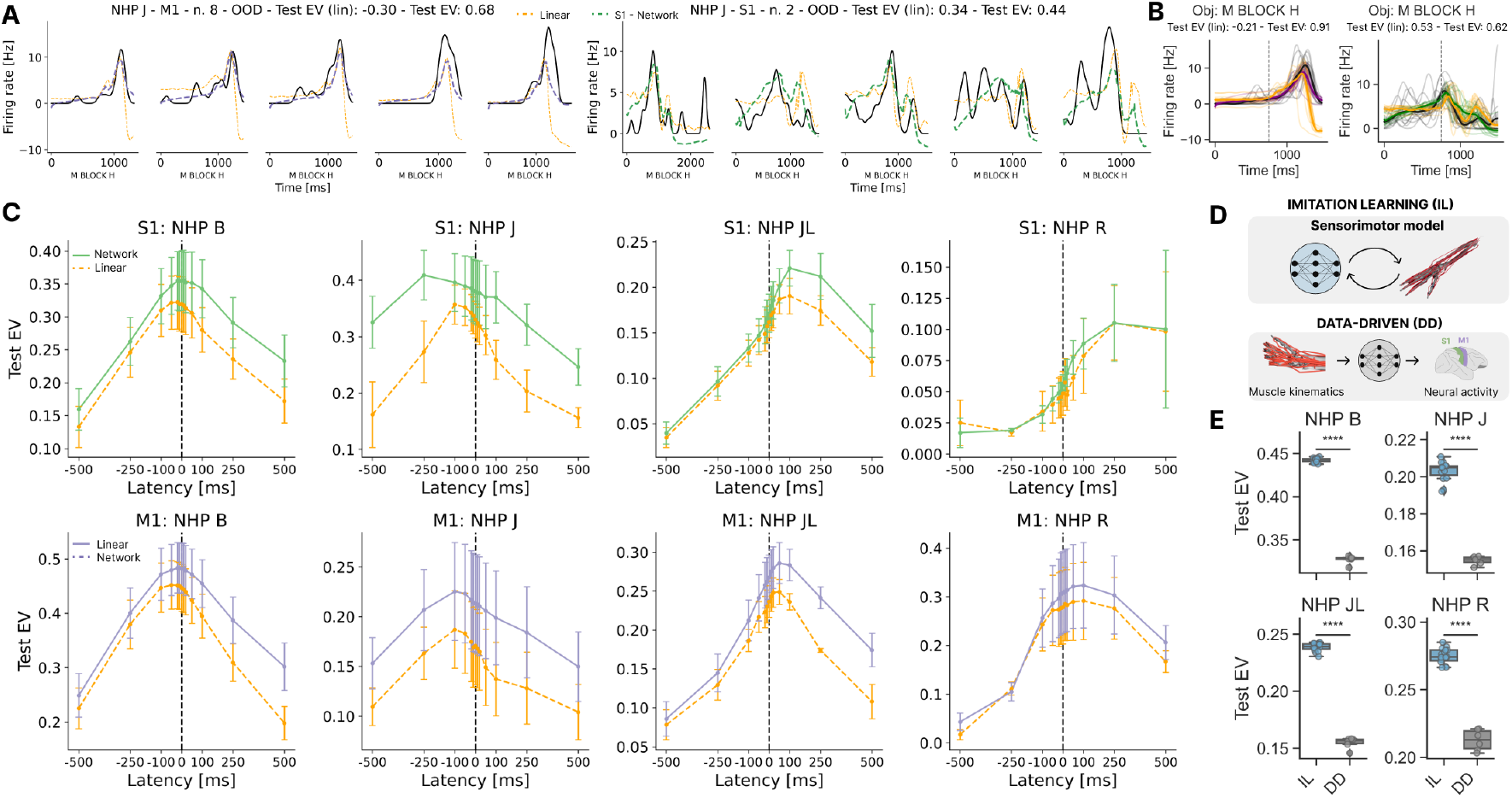
**A**. Example neural predictions on OOD test trials for a representative unit in M1 (left) and S1 (right). Black lines represent experimental spike firing rates, orange dashed lines indicate predictions from baseline linear models, and purple and green dashed lines correspond to predictions from the neural network policy. Specifically, the predictions are derived from the first MLP layer of the value network. **B**. Trial-averaged predictions for the same example neurons and objects as in **A**. Thick black lines: condition-averaged firing rates; colored lines: trial-averaged predictions; thin lines: individual trial predictions; shaded areas: s.e.m. **C**. Neural explainability comparison between the neural network policy and baseline linear models across different latencies on OOD test trials. The analysis demonstrates that neural network representations consistently outperform baseline models across all NHPs and latency conditions, highlighting the robustness of the learned representations. **D**. Schematic representation of the different frameworks used to train the neural network: Imitation Learning (IL) and Data-Driven (DD). **E**. Neural explainability distributions of imitation learning trained models agains data-driven ones. Statistical significance is evaluated using the Mann-Whitney test.

**Fig. S4.**
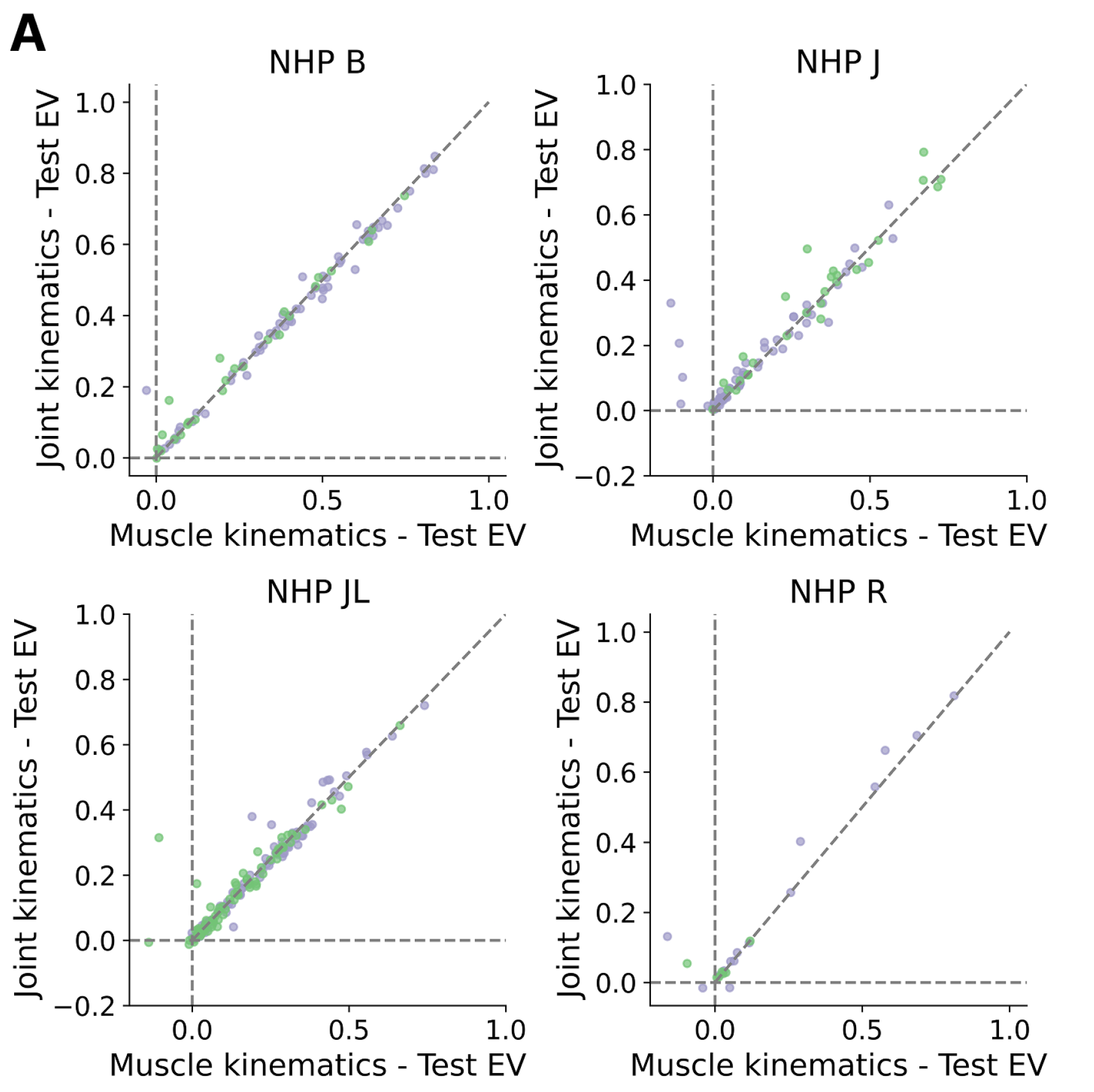
**A**. Single-neuron pairwise neural prediction comparison between joint-kinematics and muscle kinematics for individual NHP. This shows there is no predictive difference at the input level between joints and muscles.

**Fig. S5.**
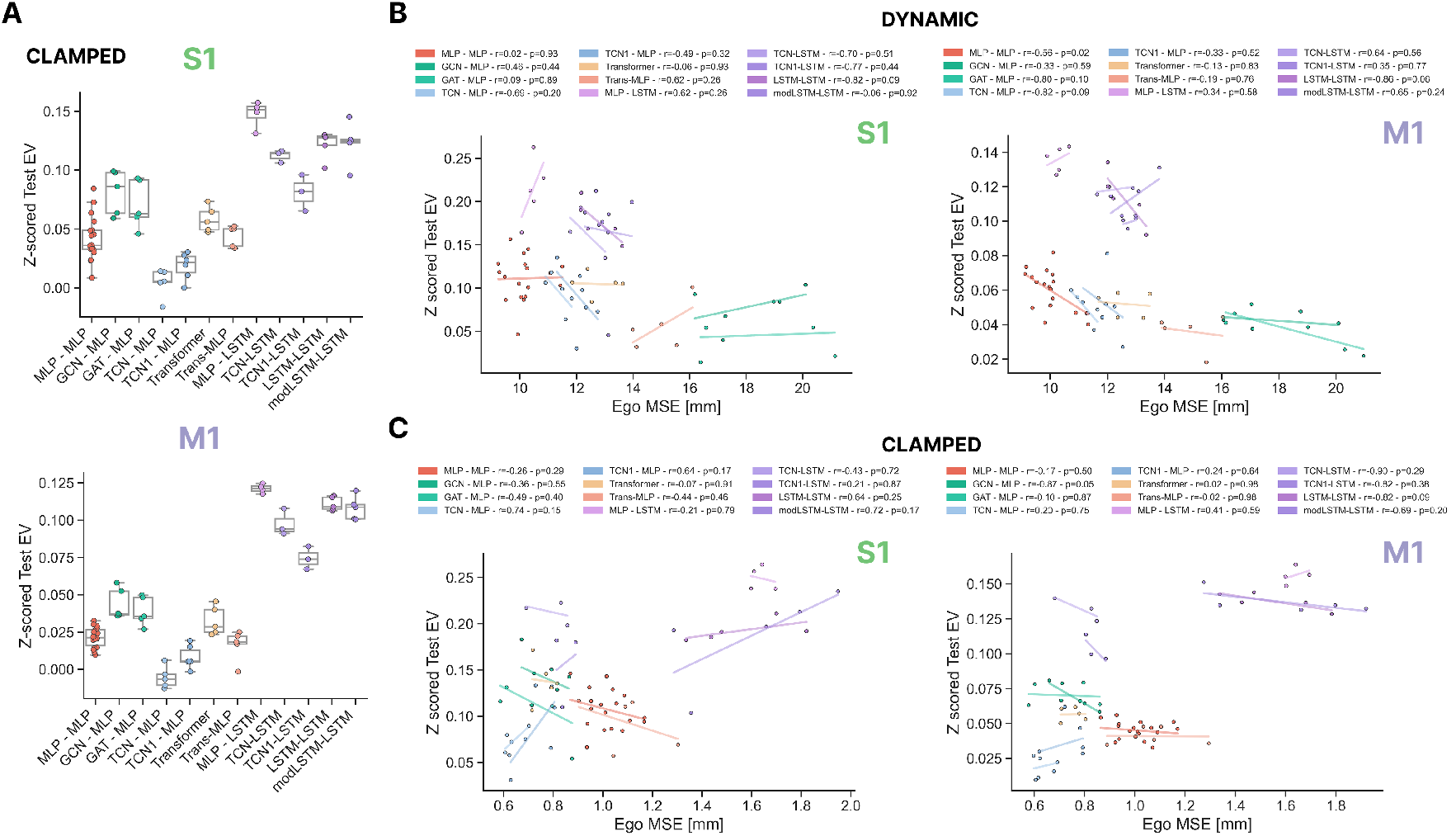
**A**. Neural alignment across architectures in the clamped-feedback condition, quantified as z-scored explained variance (EV) relative to within-animal baselines, enabling pooling across non-human primates (NHPs). Results are shown separately for S1 (top, green) and M1 (bottom, purple). Each dot represents a single trained network. Architectures with recurrent or adaptive temporal integration (LSTM, modLSTM) yield consistently higher neural predictability than feedforward or graph-based models. **B**. Relationship between neural predictability (z-scored Test EV) and behavioral performance (egocentric MSE) for the dynamic-feedback condition. Each point corresponds to one network instance. **C**. Same analysis as in **B** but under clamped-feedback conditions.

**Fig. S6.**
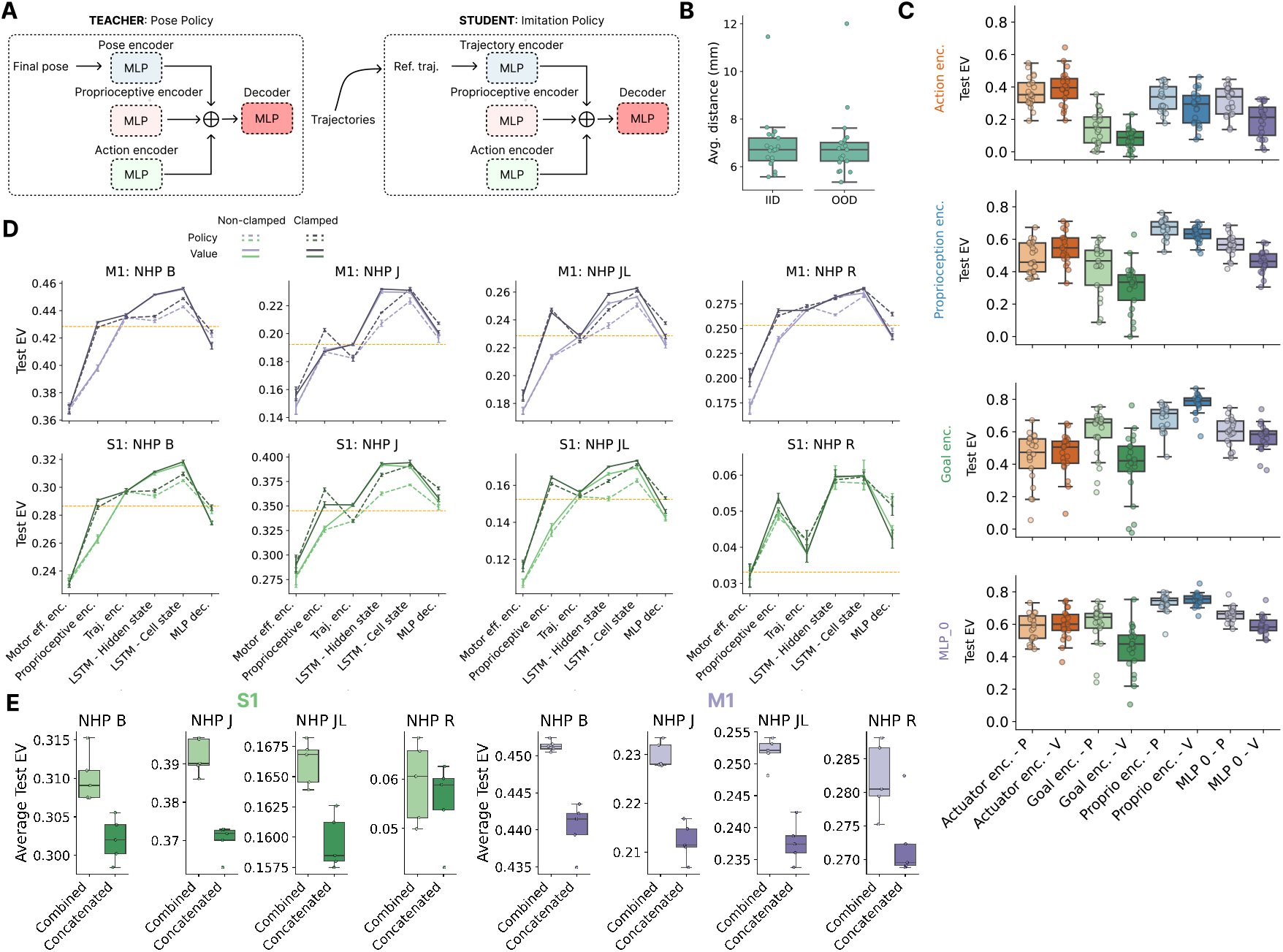
**A**. Diagram illustrating the architecture of the “Pose Policy”, trained to achieve a predefined pose, and the “Imitation Policy”, trained to reproduce the Pose Policy’s trajectory. This setup creates a synthetic control experiment for analyzing network mappings. **B**. Distribution of the average distance between the trajectories generated by the Imitation Policy and the Pose Policy, demonstrating that the Imitation Policy accurately tracks the behavior of the Pose Policy. **C**. Distribution of neural predictions across sessions, where each row represents predictions of a source layer from the Imitation Policy for a corresponding target layer in the Pose Policy. Results highlight that the encoder module of the Imitation Policy best predicts its corresponding module in the Pose Policy. However, overlapping and correlated representations make it more challenging to disentangle unique contributions, particularly in integrated modules such as the decoder. **D**. Explained variance (EV) of neural predictions across successive layers of the value (solid lines) and policy (dashed lines) networks for each NHP, shown separately for M1 (top, purple) and S1 (bottom, green). Lighter shades indicate activations extracted under natural feedback (non-clamped) conditions, and darker shades under fixed feedback (clamped) conditions. Across animals, the value network consistently outperformed the policy network, and clamped extraction enhanced alignment, suggesting that stable, state-based representations capture cortical dynamics more faithfully than raw control signals. **E**. Comparison between the integrated LSTM decoder representation (integration of proprioceptive, goal, and efferent streams) and the concatenated combination of these inputs. For both S1 and M1, the integrated representation consistently yielded higher neural predictivity, demonstrating that cortical alignment arises from the nonlinear integration of input streams rather than their simple aggregation.

**Fig. S7.**
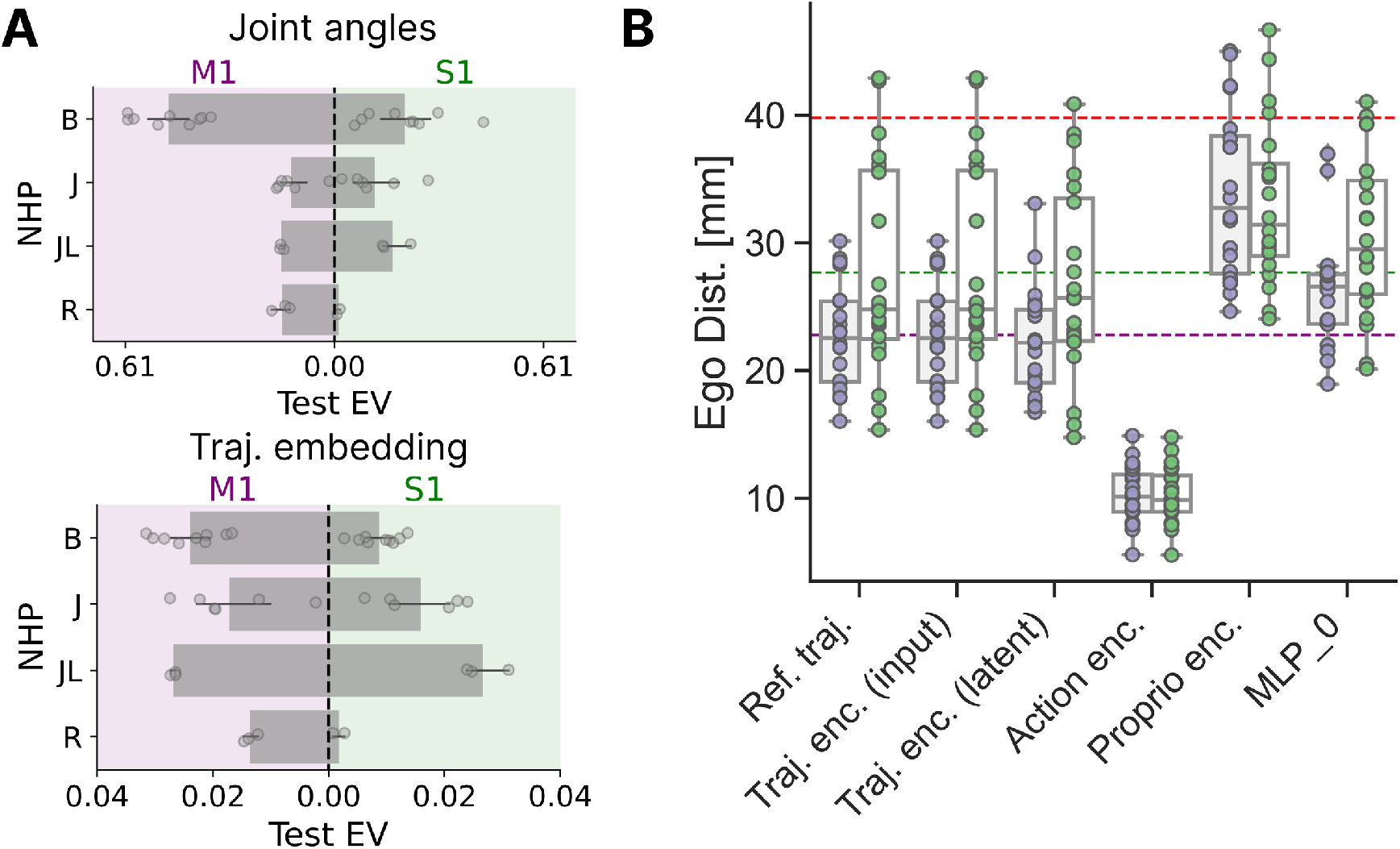
**A**. Distribution of test EV for decoding joint angles and trajectory embeddings from sensory and motor brain areas for each NHP. **B**. Behavioral performance of the *brain-controlled policy* for different decoded layers. Each dot corresponds to a session, showing the egocentric distance between the brain-controlled and reference trajectories. Dashed horizontal lines indicate baseline errors from direct joint-angle decoding (green for S1, purple for M1); the red dashed line marks the error from maintaining the average hand posture.

